# Integrated AlphaFold2 and DEER investigation of the conformational dynamics of a pH-dependent APC antiporter

**DOI:** 10.1101/2022.04.13.488180

**Authors:** Diego del Alamo, Lillian DeSousa, Rahul M. Nair, Suhaila Rahman, Jens Meiler, Hassane S. Mchaourab

## Abstract

The Amino Acid-Polyamine-Organocation transporter GadC contributes to the survival of pathogenic bacteria under extreme acid stress by exchanging extracellular glutamate for intracellular GABA. Its structure, determined exclusively in an inward-facing conformation at alkaline pH, consists of the canonical LeuT-fold of a conserved five-helix inverted repeat, thereby resembling functionally divergent transporters such as the serotonin reuptake transporter SERT and the glucose-sodium symporter transporter SGLT1. However, despite this structural similarity, it is unclear if the conformational dynamics of antiporters such as GadC follows the blueprint of these or other well-studied LeuT-fold transporters. Here, we used double electron-electron resonance (DEER) spectroscopy to monitor the conformational dynamics of GadC in lipid bilayers in response to acidification and substrate binding. To guide experimental design and facilitate the interpretation of the DEER data, we generated an ensemble of structural models in multiple conformations using a recently introduced AlphaFold2 methodology. Our experimental results reveal acid-induced conformational changes that dislodge the C-terminus from the permeation pathway coupled with rearrangement of helices that enable isomerization between both inward- and outward-facing states. The substrate glutamate, but not GABA, modulates the dynamics of an extracellular thin gate without shifting the equilibrium between inward- and outward-facing conformations. In addition to introducing an integrated methodology for probing transporter conformational dynamics, the congruence of the DEER data with patterns of structural rearrangements deduced from ensembles of AlphaFold2 models illuminate the conformational cycle of GadC underpinning transport and exposes yet another example of the divergence between the dynamics of different functional families in the LeuT-fold.

**SIGNIFICANCE STATEMENT:** The transporter GadC contributes to acid resistance in bacterial pathogens by exchanging two substrates, glutamate and GABA, using a mechanism termed alternating access. In this study, the conformational dynamics underlying alternating access was studied using a combination of spectroscopy and computational modeling. A conformationally diverse ensemble of models, generated using AlphaFold2, guided the design and interpretation of double electron-electron resonance spectroscopy experiments. We found that whereas GadC was inactive and conformationally homogeneous at neutral pH, low pH induced isomerization between two conformations. From our integrated computational/experimental investigation emerges a transport model that may be relevant to eukaryotic homologs that are involved in other cellular processes.

## INTRODUCTION

Found in all domains of life, transporters in the Amino Acid-Polyamine-Organocation (APC) family, shuttle amino acids and their derivatives across lipid bilayers^1–4^. They are presumed to mediate substrate translocation via an alternating access mechanism that entails isomerization between inward-facing and outward-facing conformations in response to substrate binding and/or release^5,6^. Substrate leak across the membrane is avoided by one of two coupling mechanisms: symport, which involves cotransport of substrates and ions in the same direction followed by substrate-free isomerization; and antiport, where alternating access is strictly ligand-dependent, and one substrate’s import is followed by the other substrate’s export. Although the APC family consists of both symporters and antiporters^3,7^, dysregulation and/or dysfunction of amino acid antiporters in humans have been shown to contribute to genetic diseases such as phenylketonuria, cystinuria, and lysinuric protein intolerance^8–16^. Upregulation of the broad-specificity amino acid transporter LAT1 (also known as SLC7A5) and/or the cystine/glutamate exchanger xCT (SLC7A11) is a hallmark of cancer that correlates with poor prognosis and survival^14–17^. LAT1 has taken further significance due to its involvement in trafficking drugs and prodrugs across the blood-brain barrier^11,17,18^.

Although prokaryotes rely on APC transporters to acquire and retain amino acids, the most well-studied antiporters are those used by bacterial pathogens such as *E. coli* O157:H7 to withstand extreme acid stress^19–24^. These include the glutamate/GABA antiporter GadC and the arginine/agmatine antiporter AdiC, which import the precursors and export the products of proton-consuming amino acid decarboxylases responsible for raising intracellular pH^23–25^. AdiC adopts a homodimeric assembly, while GadC possesses a C-terminal tail that putatively regulates pH-dependent transport^26–28^ (Figure 1A). Nevertheless, despite these unique adaptations and their functional divergence from human transporters, both exchangers have served as model systems for eukaryotic homologs since their initial structural characterization over a decade ago^29–32^. The crystal structures of GadC and AdiC, which were determined in inward-facing and outward-facing conformations, respectively, superimpose over recent cryo-EM structures of human orthologs and have been used, for example, for numerous drug docking studies^33–38^. Nevertheless, an absence of detailed studies of their energy landscapes and conformational dynamics has hindered framing these structures in the broader context of antiport.

**Figure 1.**
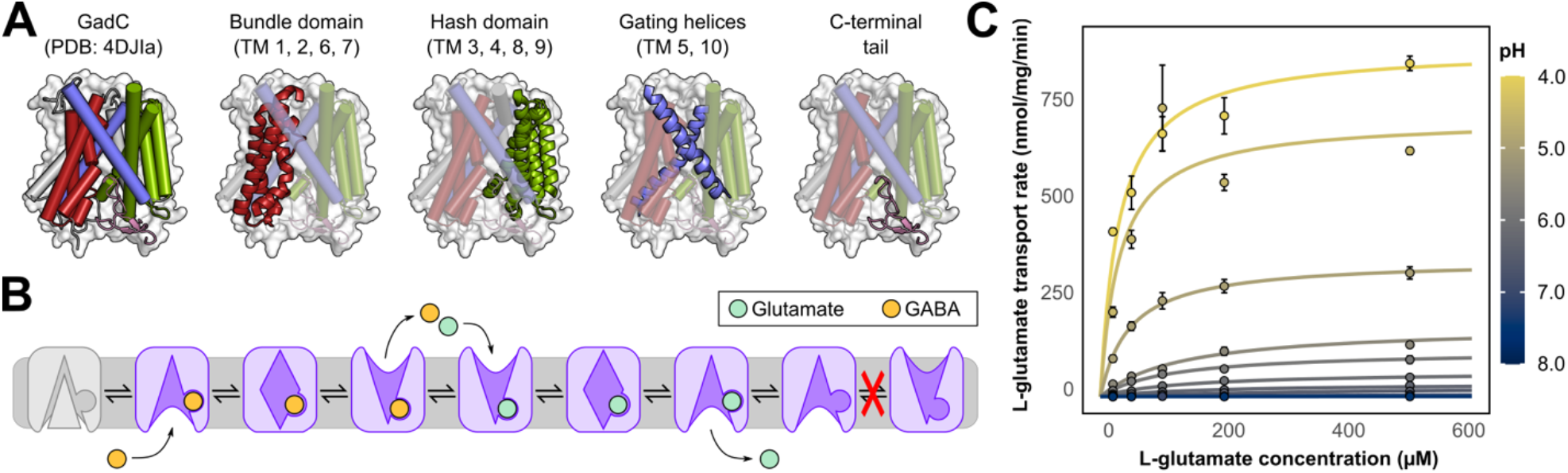
Architecture and pH-dependent transport activity of GadC. **(A)** The crystal structure of GadC consists of a four-helix bundle domain, a four-helix hash domain, two gating helices, and two ancillary helices uninvolved in transport (not emphasized). Additionally, the C-terminal tail is wedged into the intracellular cavity, locking the transporter into a putatively inactive conformation. **(B)** Proposed antiport mechanism consisting of inward-facing, outward-facing, and occluded conformations. This mechanism implicates glutamate and GABA in driving the outward-to-inward and inward-to-outward conformational transition, respectively, and it forbids substrate-free isomerization. **(C)** Glutamate transport by wildtype GadC reconstituted into proteoliposomes is pH-dependent. Error bars correspond to the standard error of the mean (n=3).

Insights in this regard may be obtained from more distant homologs of APC transporters that have been subjects of extensive study^39–51^. These include the neurotransmitter-sodium symporter SERT^52,53^, the sodium-solute symporter SGLT1^54^, and bacterial model systems such as the leucine transporter LeuT^55,56^, the benzylhydantoin transporter Mhp1^57,58^, and the betaine transporter BetP^59,60^. A broad record of experimental investigation into the conformational dynamics demonstrates elements that are both conserved (such as substrate-dependent isomerization) and divergent (such as the effect of sodium binding on conformational dynamics)^61^. However, it is unclear if patterns derived from the study of these LeuT-fold symporters extend to antiporters, particularly since the former must undergo a ligand-free isomerization step that is inhibited in the latter^5^ (Figure 1B).

Here we report an investigation of the conformational dynamics of GadC using double electron-electron resonance (DEER) spectroscopy^62^, a technique that has been successfully applied to the study of LeuT-fold symporters such as LeuT and Mhp1^39–41,63^. To facilitate the interpretation of these results, we capitalized on our recent modification of the structure prediction algorithm AlphaFold2^64,65^ (AF2) to generate structural models of GadC in multiple conformations. DEER distance distributions between spin label pairs designed based on these models suggest that the structure of GadC at acidic pH is substantially more dynamic than at neutral pH. Moreover, the C-terminal tail, putatively responsible for regulating transport at neutral pH, detaches and becomes disordered under weakly acidic conditions. At low pH, GadC is in conformational equilibrium between inward-open and outward-open conformations, with experimental distance changes in agreement with predictions made from the AF2 models relative to the crystal structure. Substrate binding did not shift this equilibrium between conformations beyond an extracellular thin gate, a finding which contrasts with a panel of previously investigated symporters^39,40^. This substrate-specific occlusion of the extracellular vestibule may be relevant to eukaryotic homologs such as xCT, which exchange amino acids in different directions with high specificity^14^. Moreover, because similar elements of alternating access were observed in the sodium-hydantoin symporter Mhp1, commonalities may exist in the transport cycles of these distantly related transporters.

## RESULTS

The pH-dependent activity profile of GadC was verified by measuring radiolabeled substrate uptake into proteoliposomes. A construct of wildtype GadC cloned from *E. coli* str. O157:H7 was expressed in *E. coli* C43 (DE3), purified in β-DDM detergent micelles, and reconstituted into proteoliposomes containing 5 mM glutamate at pH 5.5 (see *Materials and Methods*). These proteoliposomes were then tested for substrate transport by detection of [^3^H]-L-glutamate uptake as a function of both external pH and substrate concentration. Additionally, time-dependent glutamate transport was measured in proteoliposomes containing 5 mM GABA at pH 5.5 (Figure S1). Consistent with previous findings^26,66^, we observed a strong dependence of radioligand uptake on pH: increasing the pH from 4.0 to 6.5 reduced glutamate transport by about 97% (Figure 1C).

To characterize the structural changes associated with pH-dependent activation of transport, we used site-directed spin labeling and EPR spectroscopy^62,67^. All three endogenous cysteines in the wildtype sequence were mutated to chemically inert residues (C60V, C247A, C380V). As with previous studies on structural homologs of GadC, double-cysteine pairs were selected based on their ability to report on inter- and intra-domain movements. To evaluate if these measurements were expected to fall within the detectable range for DEER measurements (15-60 Å) and to test whether the resulting data were consistent with the crystal structure, distance measurements were first simulated between candidate residue pairs using dummy spin labels explicitly modeled on the crystal structure^68,69^. Following purification and spin-labeling, all mutants were reconstituted into proteoliposomes and tested for transport and pH-dependent inactivation at pH 5.5 (Figure S2) and 7.5 (Figure S3), respectively. Additionally, all experimental DEER measurements were carried out in nanodiscs composed of the same lipids as those of the proteoliposomes used for transport assays. This ensured that neither the spin labels nor the membrane environment interfered with GadC’s ability to traffic substrates at acidic pH or to undergo inactivation at neutral pH. Finally, we note that although GadC was crystallized as an antiparallel homodimer, no experimental evidence of this quaternary assembly was detected in lipid nanodiscs.

### pH-dependent detachment of the C-terminus is required for activation

Abrogation of transport at neutral and alkaline pH has previously been attributed to the transporter’s C-terminus (residues 471-511, shown in pink in Figures 1A and 2A). In the crystal structure of GadC, captured at pH 8.0, the C-terminal tail is embedded in the intracellular cavity and obstructs both substrate passage and closure of the intracellular gate, two prerequisites of alternating access^26^. To test the hypothesis that this domain detaches under acidic conditions, a double-cysteine mutant (L143C/E480C) was generated to measure the distance between spin labels at the C-terminus and the transmembrane domain. Distance distributions of this pair reported large changes as a function of pH. At neutral pH, the average distance matched that predicted from the crystal structure, whereas a sharp increase in the magnitude, distance, and the width of the distribution was observed under acidic conditions (Figures 2B and S4). A nonlinear least-squares fit to the amplitudes of the short- and long-distance components revealed that this shift occurred cooperatively with a pKa of 6.07±0.11 (Figure 2C), with both short-distance components diminishing at low pH in a tightly correlated manner (Figure S5).

**Figure 2.**
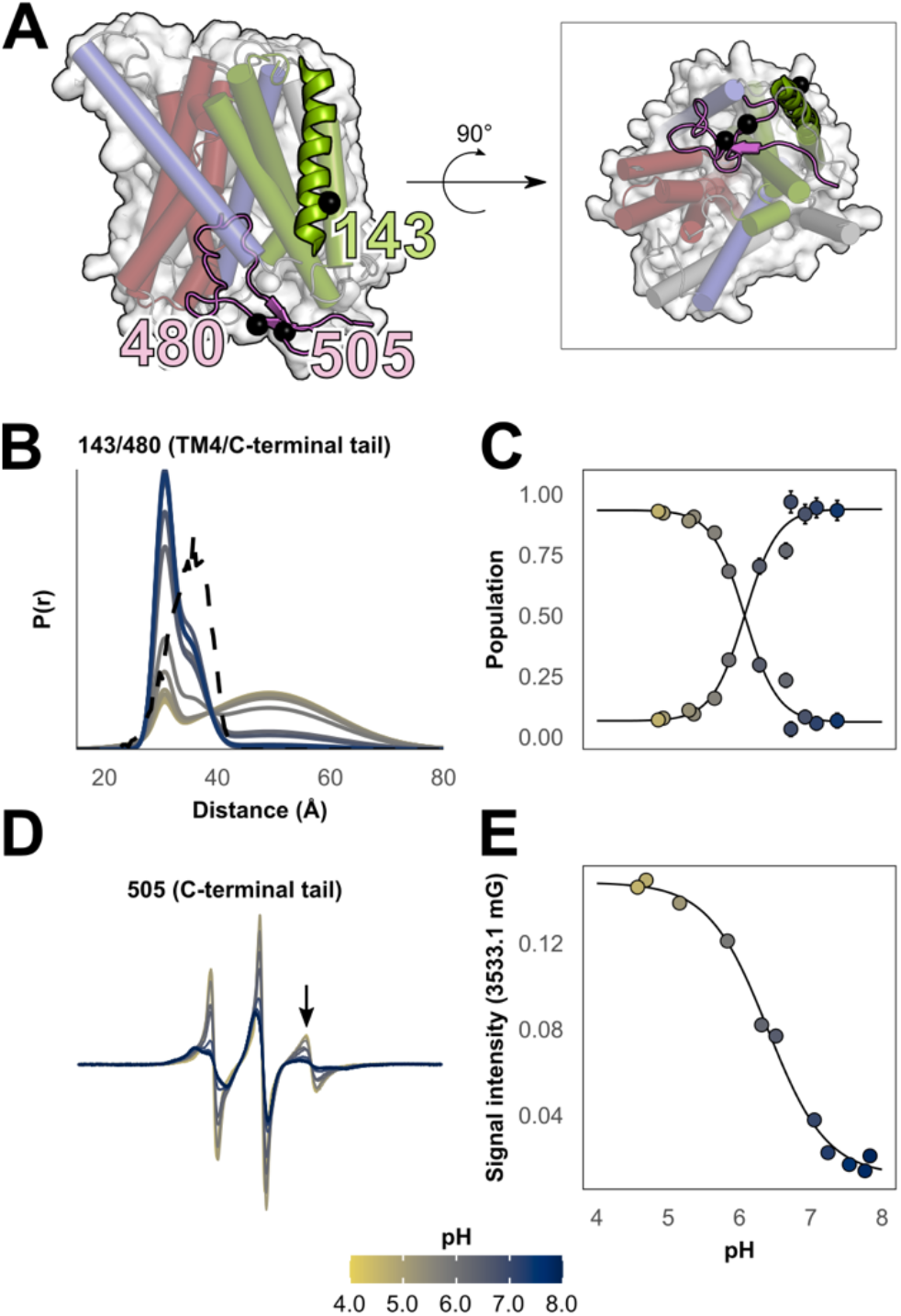
Detachment of the C-terminus of GadC is triggered by low pH. **(A)** Position of the C-terminal tail, shown in pink, relative to the main transmembrane domain of the transporter. Inset: rotated view of the C-terminal tail embedded in the intracellular vestibule. **(B)** At low pH, a broad long-distance component is observed in equilibrium with a distance component consistent with predictions made from the crystal structure (shown in the dashed line). **(C)** Titration measurement of the dissociation of the C-terminus. Error bars correspond to 95% confidence intervals calculated using the program GLADDvu (see *Materials and Methods*). **(D)** pH-dependent increases in conformational heterogeneity resolved by continuous-wave EPR. **(E)** Titration measurement of the mobile component of the CW spectra reveals a similar profile to the DEER measurements.

To further determine if this cooperative distance change originated from conformational disorder of the C-terminal domain, a single-cysteine mutant was introduced (V505C). At neutral pH, the lineshape of this spin-labeled mutant’s continuous wave EPR spectrum suggested that the domain was relatively structured, consistent with its docked conformation in the crystal structure (Figure 2D). By contrast, reducing the pH led to a sharp, highly mobile spectral component that dominated the lineshape at pH 6.0 and below, suggesting an increase in structural disorder. Nonlinear least squares fit of the EPR spectrum high field line as a function of pH yielded a pKa of 6.30±0.04, consistent with the DEER measurements discussed above (Figures 2D and 2E). Taken together, the data corroborate the hypothesis that the tail detaches from the transmembrane domain and becomes heterogeneous and disordered. Additionally, the pKa of this event closely matched the pH at which transport activity is abrogated, reinforcing this domain’s role in regulating substrate exchange under neutral pH conditions.

### Modeling alternative conformations of GadC using AlphaFold2 and Rosetta

Next, we sought to characterize the movements underpinning the transport cycle of GadC using a library of double-cysteine mutants selected to detect pH- and ligand-dependent structural changes. An outward-facing homology model, generated with RosettaCM^70^ using structures of the arginine/agmatine transporter AdiC as templates^27,28,71,72^, initially guided experimental design but was discarded when predicted distance changes failed to correspond to experimental DEER results (Figure S6). We therefore used the deep learning algorithm AF2^64^ to predict the structure of GadC in multiple conformations following detachment of the C-terminus by introducing several modifications described in detail in *Materials and Methods* and elsewhere^65^. Model generation was followed by constrained refinement in an implicit membrane using Rosetta^73–75^. To identify the breadth of the energy minimum with greater confidence, we generated 650 models and visualized the predicted conformational changes using dimensionality reduction with principal component analysis (PCA, Figure 3).

**Figure 3.**
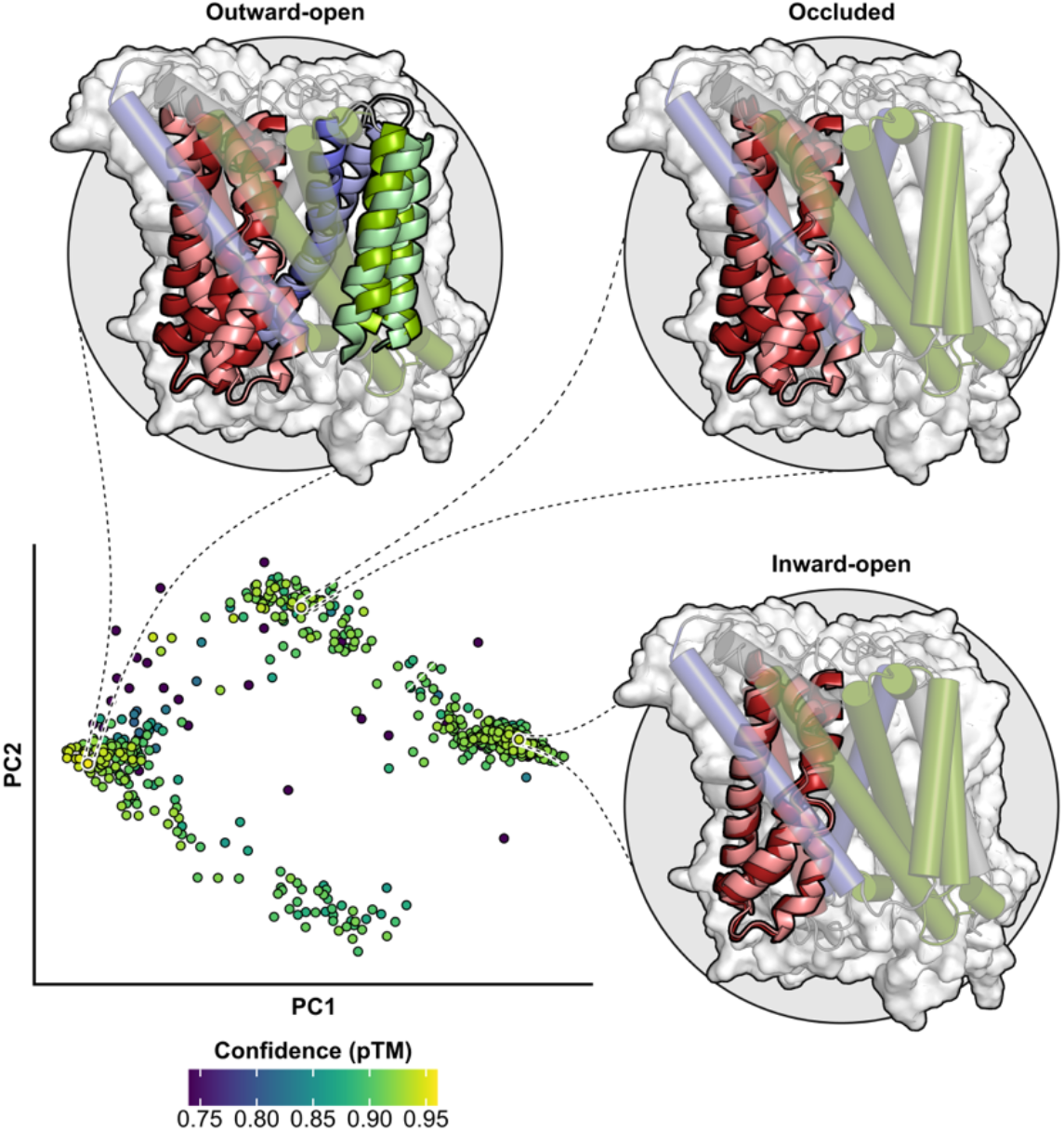
Alternative conformations of GadC modeled by AlphaFold2. Dimensionality reduction using PCA (bottom left) revealed three densely populated clusters with inward-open, fully occluded, and outward-open models. Helices predicted to move relative to the crystal structure are shown as ribbons, with darker ribbons corresponding to those of the inward-facing crystal structure and lighter ribbons to the respective AF2 models. The highest-confidence models of each cluster in the PCA plot are shown above in light colors (model confidence was measured using pTM, with greater values indicating greater model confidence). A fourth cluster with doubly-open models (bottom center) was not analyzed.

Models generated with high confidence were distributed across three clusters, and a substantial fraction were dissimilar to the crystal structure. Visual inspection suggested that members of these three clusters could be assigned to either inward-open, fully occluded, and outward-open conformations (a fourth cluster, which consisted of doubly-open models, was sparsely populated and therefore ignored during our analysis). The inward-open models differed from the crystal structure primarily in transmembrane helices 1, 6, and 7, which become slightly pinched on the intracellular side. The occluded models suggested that closure of the intracellular cavity was mediated by a large-amplitude movement of these same helices. Finally, opening of the extracellular side in the outward-open models involved translations by TM1 and TM6, rotation of TM4 and TM9 in the hash domain, and straightening of TM10. Overall, the proposed mechanism closely resembles that of the homolog LAT1 while deviating from that of AdiC^28,76,77^.

A representative model from each cluster was selected based on predicted confidence values (pTM). As with the crystal structure, distance distributions between spin labels were predicted from these models for comparison to experimental DEER measurements. With these models in hand, we set out to design double-cysteine pairs to test the implied structural changes and determine the conditions under which they are sampled. For clarity, the following Figures show only the OF model; predictions made from the IF and occluded models are shown in Figure S7.

### Evaluation of the rocking bundle model of alternating access

The extent to which the bundle domain, which comprises transmembrane helices 1, 2, 6, and 7, undergoes a rigid-body conformational change was assessed using a two-pronged approach. First, a network of double-cysteine mutants, spanning both sides of the membrane, probed the magnitude of the movements undertaken by these helices relative to reference residues located on helices not predicted to move (Figures 4 and 5). Second, intra-bundle measurements determined whether the domain itself isomerizes as a rigid body (Figure 6). Whereas this movement was initially proposed based on comparisons of the crystal structure to outward-facing structures of AdiC^26^, the ensemble of AF2 models instead predicts substantial bending and independent movement of bundle domain helices.

**Figure 4.**
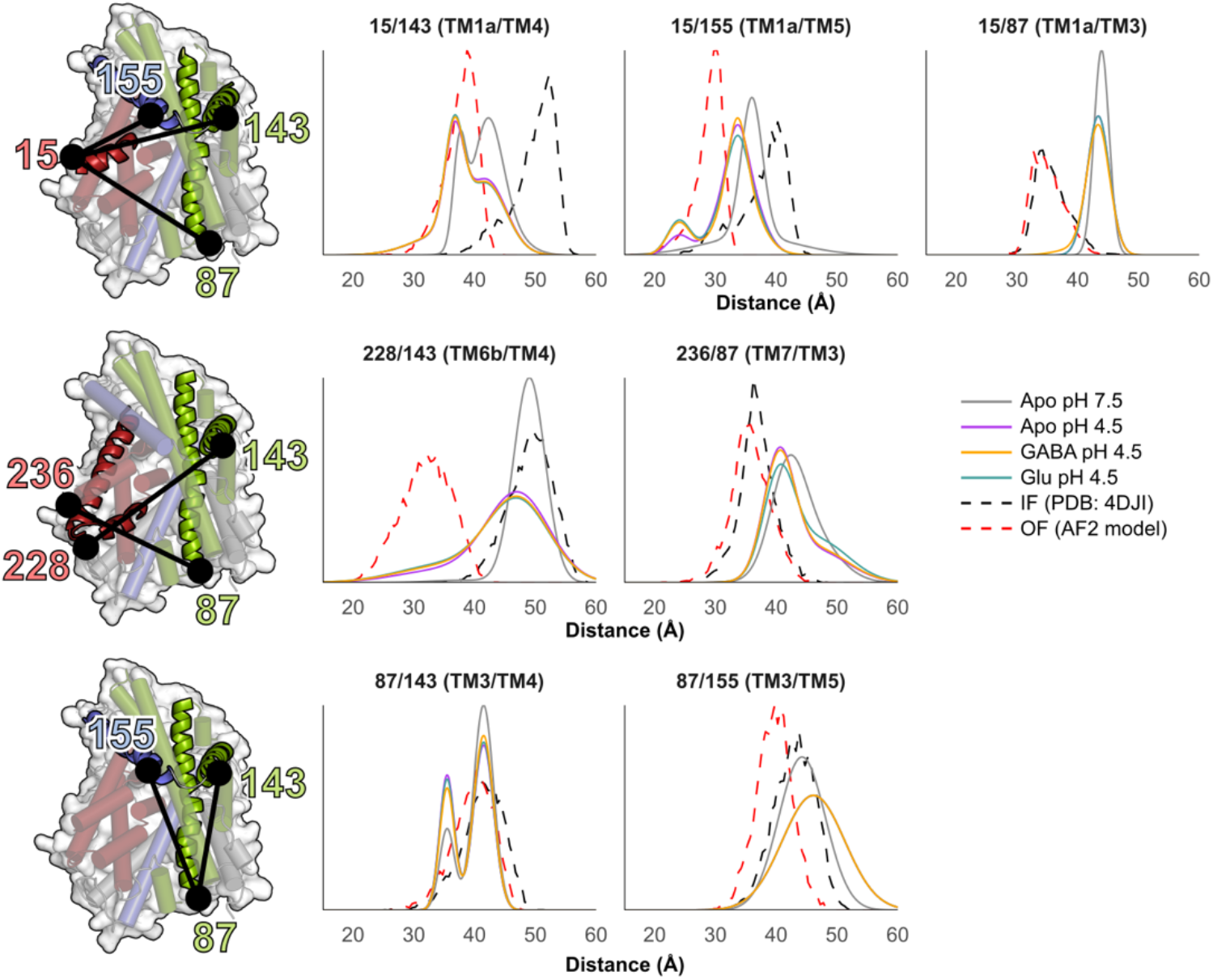
Conformational changes in the bundle domain on the intracellular side. Cartoon depictions of GadC shown from the intracellular side. Labeled positions are indicated as black spheres in cartoons, with the bundle domain shown in red. Distance distributions between the bundle domain and reference sites (in blue and green) are largely unimodal at neutral pH and become more heterogeneous at low pH. Short-distance components consistent with an inward-closed model were also observed in some distributions. Substrates were not observed to affect the distance distributions.

**Figure 5.**
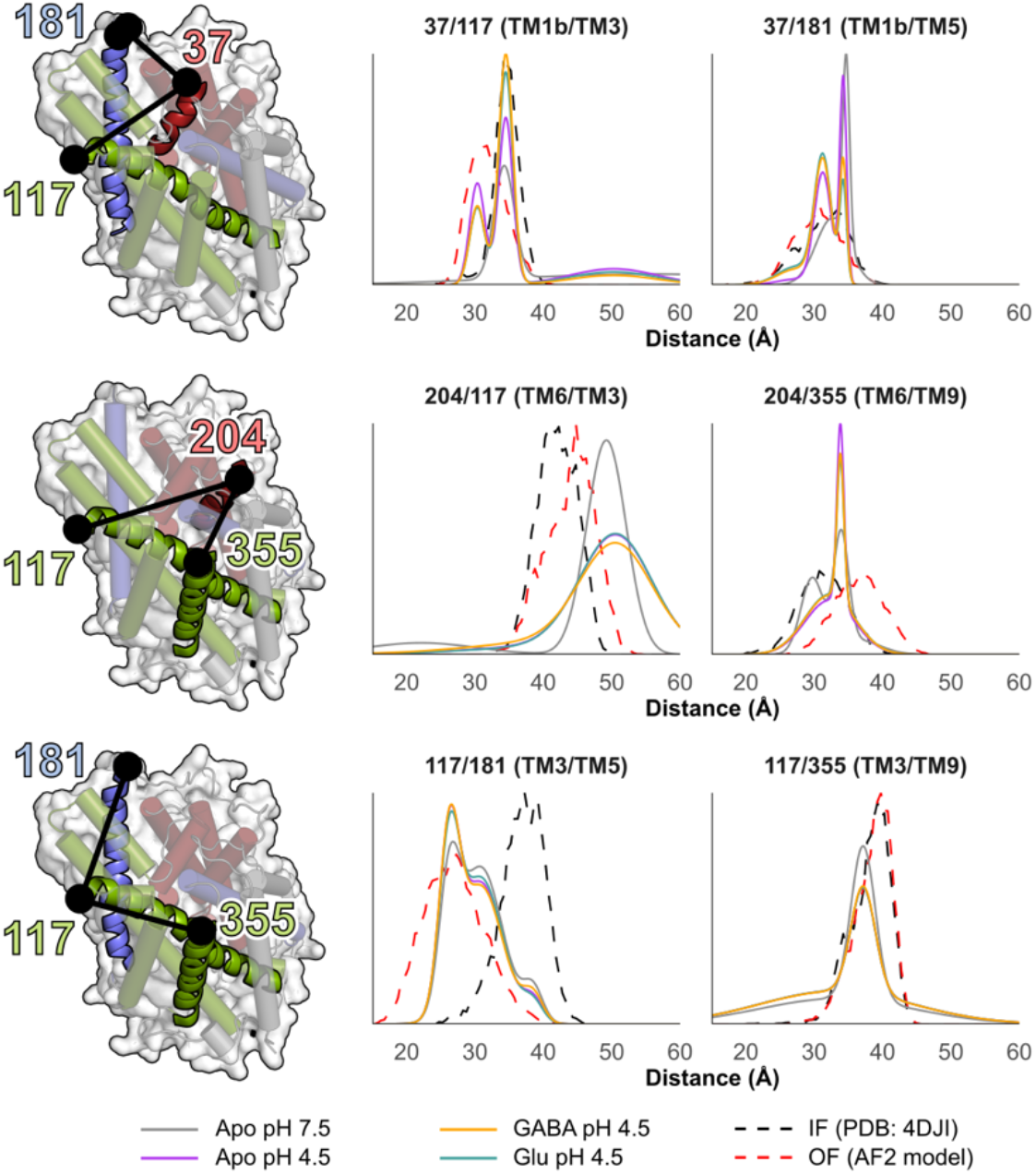
Conformational changes in the bundle domain on the extracellular side. Cartoon depictions of GadC shown from the extracellular side, with labeled positions indicated as black spheres. Distance distributions overlapped with predictions made from the crystal structure at neutral pH. Low pH coincided with equilibrium shifts toward populations consistent with an outward-open model in a substrate-independent manner.

**Figure 6.**
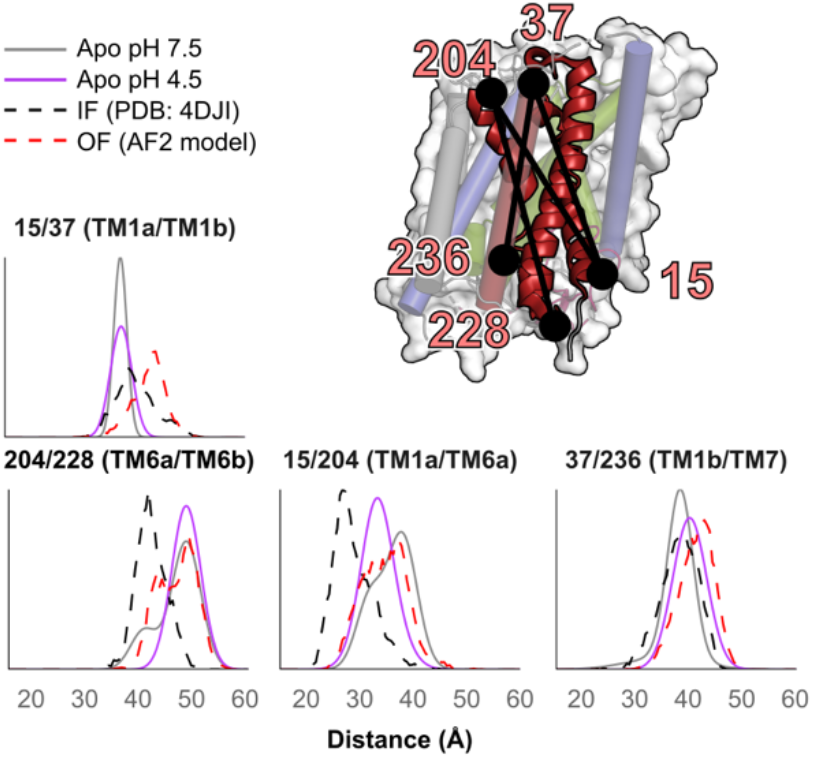
The bundle domain of GadC does not behave as a rigid body. Measurements between spin labels (indicated by black spheres) across the membrane indicate a degree of intra- and inter-helical flexibility. Low-pH measurements with GABA and glutamate overlapped with Apo low-pH measurements and are omitted for clarity.

Distance measurements carried out at neutral pH without substrate indicated a largely homogeneous population (Figures 4, 5, S9, and S10). Unimodal peaks were observed on the intracellular side of the membrane at neutral pH, and experimental distributions closely aligned with predictions made from the crystal structure on the extracellular side.

Decreasing the pH to 4.5, which falls below the pKa of C-terminal detachment, induced more heterogeneity on the intracellular side. Distributions between spin label pairs involving TM1 broadened and/or became multicomponent, indicating a degree of backbone flexibility absent at neutral pH. These results indicated movement of TM1 and TM6 into the intracellular cavity to partially replace the space vacated by the C-terminal tail (Figure 3). Additionally, lower-amplitude short-distance peaks and/or shoulders were observed in TM1/TM4, TM1/TM5, and TM4/TM6, consistent with the inward-closed AF2 models. Notable deviations between predicted and experimental distributions involving residues 15 (TM1a) and 87 (TM3) were observed, possibly due to their locations near crystal contacts (Figure S14).

On the extracellular side, measurements involving TM1 were also consistent with partial adoption of an outward-open conformation at low pH. Distance changes involving TM6, meanwhile, also appeared to indicate a partial population shift toward conformations similar to the outward-open models predicted by AF2. Overall, the data indicate that, at low pH, the bundle-domain moves into the intracellular cavity vacated by the C-terminal domain following detachment as predicted by the inward-open AF2 model. They also suggest the concomitant sampling of an outward-open conformation consistent with the AF2 model.

A notable finding was the lack of glutamate- or GABA-dependent conformational dynamics evident in either the DEER distributions or the CW spectra in the bundle domain. While very slight increases in inward- and outward-closed populations were observed, the confidence intervals of these changes overlapped with those of GadC under apo conditions (Figures S9 and S10). We therefore concluded that saturating concentrations of either glutamate or GABA have no impact on the conformational equilibrium of GadC in this region at low pH. This pattern recurred throughout the structure, with one notable exception discussed below. Thus, for clarity, distributions collected with either glutamate or GABA will be omitted in the text below.

### The bundle domain does not move as a rigid body

Comparison of the GadC crystal structure to that of the homolog AdiC prompted the hypothesis that the bundle domain facilitates alternating access by a rigid-body movement^26^. In contrast, the occluded and outward-facing models generated using AF2 suggest that closure of the intracellular side is facilitated by bending of TM7 and translation of TM1 and TM6 into the space vacated by the C-terminal domain. Both models were tested by measuring distributions within the bundle domain. However, the close distances of helices within the domain, combined with the 15 Å lower distance limit of the DEER technique, required that these measurements be carried out across the membrane (i.e., inner to outer leaflet).

Measurements between spin labels at either end of TM1 and TM6 show pH-dependent movement inconsistent with a rigid-body hypothesis. In structurally homologous LeuT-fold transporters undergoing rigid-body conformational changes, little to no movement is observed within the bundle domain. In contrast, decreases in pH were sufficient to induce changes in both the distance distribution (Figure 4C) and the CW spectra (Figure S11). Thus, we concluded that alternating access in GadC departs from a rigid-body motions in the bundle domain as initially proposed. Instead, this conformational change appears to incorporate elements of helical translations that have been experimentally observed by crystallography and cryo-EM in closely related homologs such as LAT1 as well as more distant homologs such as LeuT and SERT^33,52,53,55,56,77.^

### Lack of large-scale movement in IL1 or EL4

Two additional hypotheses, implied by observations in homologous LeuT-Fold transporters, propose a role in facilitating alternating access for the amphipathic loops IL1 and EL4, which are adjacent in sequence to the bundle domain on the intracellular and extracellular sides, respectively. In AdiC, a partially conserved tyrosine residue on IL1 has been experimentally shown to regulate pH-dependent transport^72^. Its mutagenesis to alanine, but not phenylalanine, abolished transport inactivation under neutral conditions. As the corresponding residue in GadC is phenylalanine, this finding hinted at a possible role in mediating pH-dependent isomerization. Therefore, we mutated the adjacent residue, Ala77, to cysteine to both probe any pH-dependent conformational changes in this domain and to observe changes to the CW spectra indicative of environmental changes. Experimental measurements between this domain and TM3 and/or TM5 ruled out any pH-dependent helical movement (Figure 7), while CW spectra argued against changes in the local environment of Ala77 (Figure S12). These results suggest that this domain is not involved in the activation mechanism of GadC and remains firmly stapled to TM3, consistent with its conservation between structures of homologous transporters^28,76,78^.

**Figure 7.**
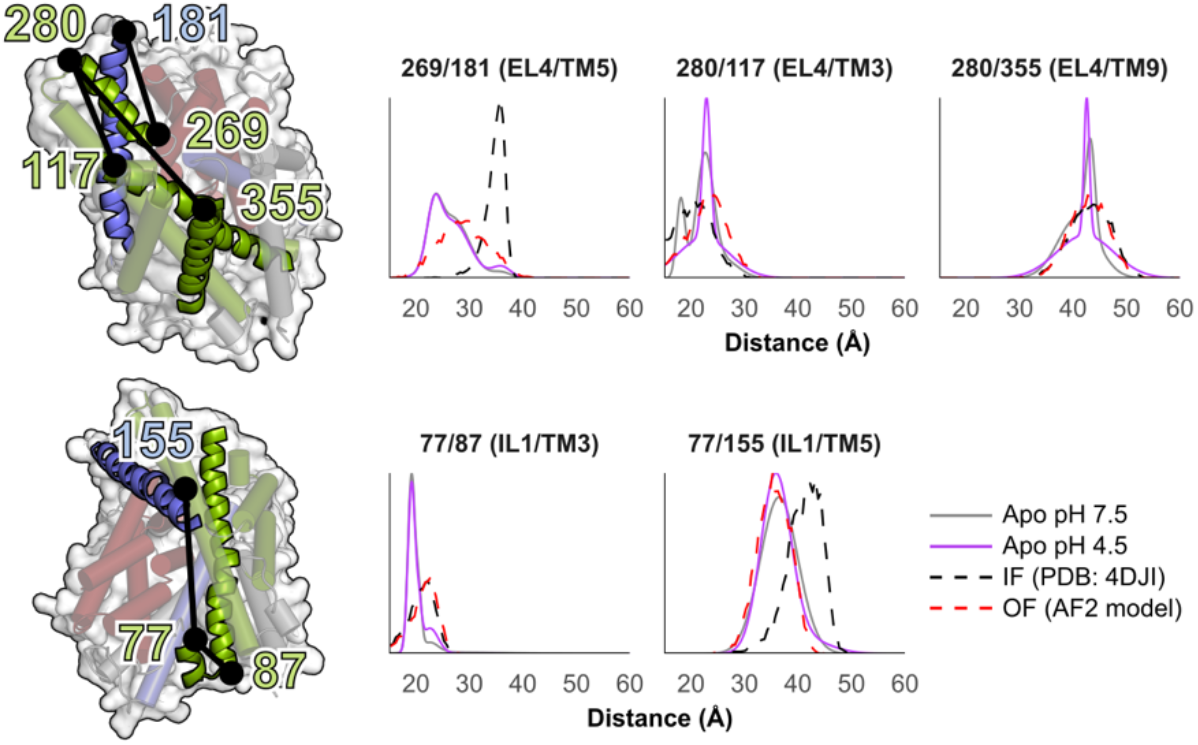
No large-scale pH-dependent changes in the positions of IL1 and EL4. Top: Position of EL4 as measured from the top and bottom of the helix (residues 269 and 280, respectively). Bottom: IL1. Cartoon depictions of GadC shown on the left along with positions of labeling sites indicated by black spheres.

Similarly, movement of EL4 has been shown in AdiC and other homologs such as sodium-coupled symporters such as LeuT, to enable access to the extracellular vestibule^28,63,76^. However, the ensemble of AF2 models did not predict an equivalent mechanism for GadC. These distinct models were tested by labeling both ends of EL4 and measuring distances to the hash domain and to TM5. Consistent with the AF2 predictions and unlike the movements observed in other LeuT-fold homologs, we observed no evidence indicating conformational changes as a function of either pH (Figure 7) or substrate addition (Figure S11) in this region. This suggests that substrate entry and exit on the extracellular side is facilitated by a different region of the transporter.

### An extracellular thin gate discriminates between GABA and glutamate

An alternative extracellular permeation pathway to that involving EL4, gleaned from the AF2 ensemble, primarily involves bending and straightening of TM10. Such a mechanism has been observed in both experimental measurements and atomistic simulations in homologous symporters such as vSGLT and Mhp1, where TM10 forms a thin gate to the substrate binding site and undergoes movement largely uncoupled from adjacent helices^39,41,57,58,79–82^. The AF2 ensemble, by contrast, predicted that this helix’s position is tightly coupled to that of TM9 in the hash domain. Distance measurements from this helix at neutral pH were defined by a single sharp component, which corroborated observations made throughout the structure (Figures 8A and B). Decreasing the pH led to a bimodal distribution with peaks spaced 10 Å apart, consistent with predictions made from comparison of the crystal structure and OF AF2 model. Notably, whereas this distribution was unchanged following addition of saturating concentrations of GABA, glutamate appeared to promote closure of this thin gate, a finding that was corroborated across multiple biological repeats (Figure S13). No such effect was observed in measurements involving the extracellular side of TM9, which is linked to TM10 by a triple-glycine motif. Thus, these data demonstrate that the dynamics of these two helices are uncoupled in the outward-facing state. Moreover, they indicate a mechanism by which GadC differentiates between its two substrates that may be relevant to homologous APC transporters.

**Figure 8.**
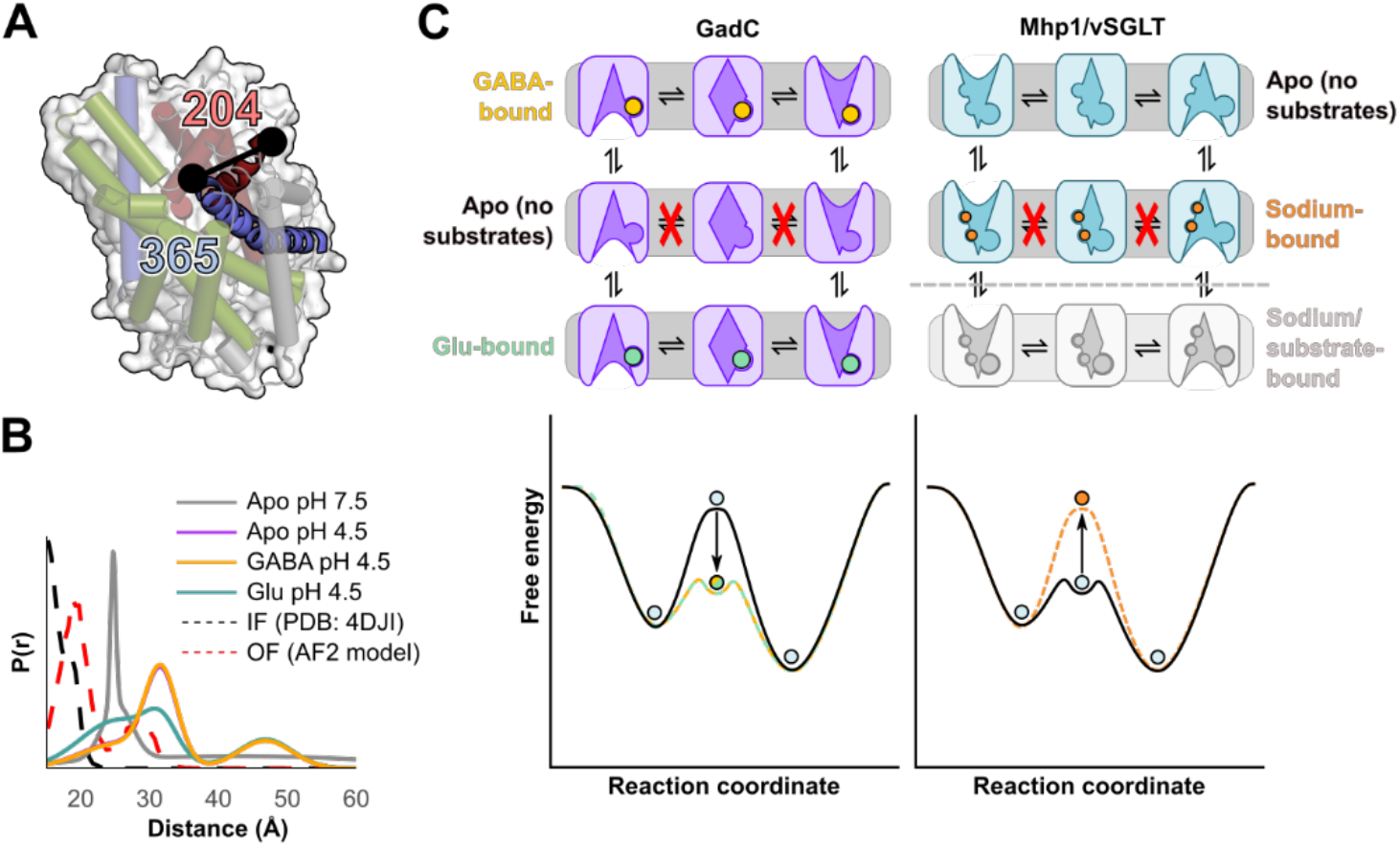
TM10 undergoes substrate-dependent dynamics independent from the hash domain. **(A)** Cartoon depiction of GadC highlighting TMs 6a and 10 and the spin labeling sites (black spheres). **(B)** Distance distributions between TM6a and TM10 demonstrate that glutamate, but not GABA (purple and yellow traces are superimposable), induces a population shift (the long-distance peak at 48 Å is likely an aggregation artifact also present in the apo pH 7.5 trace). **(C)** Proposed activation and glutamate-GABA antiport mechanism (left). The C-terminal domain detaches at weakly acidic pH, allowing isomerization to proceed. Glutamate and GABA lower an energy barrier separating the inward- and outward-facing conformations. No large-scale substrate-dependent conformational changes were observed in GadC that were equivalent to binding of sodium and substrates to sodium-coupled symporters such as Mhp1 and vSGLT (shown on the right).

## DISCUSSION

In this report, we investigated the structure and dynamics of the pH dependent Glu/GABA antiporter GadC, a member of the APC family and a homolog of human transporters implicated in various diseases, by integrating DEER spectroscopy with accurate structure prediction using AF2. This methodology, which capitalizes on a recent modification of AF2 by our group^65^, is general and is predicted to increase the throughput of spectroscopic and biochemical investigations by enabling optimal experimental design.

GadC’s conformational cycle underpins coupled glutamate import and GABA export, whereas isomerization of the substrate-free transporter is inhibited. GadC is both functionally coupled to and co-transcribed with the soluble enzyme GadB, which irreversibly decarboxylates glutamate into GABA^83^. In some species of bacteria, the intracellular concentration of GABA has been shown to reach up to 80 mM during acid stress, almost certainly exceeding that of the extracellular milieu by several orders of magnitude^84^. The differences between the concentrations of glutamate on either side of the membrane may not be as extreme^85,86^. Thus, we propose that the export step likely drives the conformational cycle of GadC.

This functional context is the foundation for one of this study’s key conclusions, i.e. the absence of detectable substrate-induced conformational changes throughout most of the structure of GadC. By contrast, pH changes shifted the equilibrium between IF and OF conformations, particularly on the intracellular side and the C-terminal tail. Indeed, substrate-induced movements were limited to the extracellular half of TM10, which was predicted by our ensemble of AF2 models to bend and straighten in the outward-closed and outward-open conformations, respectively. Glutamate, but not GABA, pushed the conformational equilibrium toward the former relative to apo conditions. However, the remainder of the structure appeared insensitive to the addition of either substrate. Thus, we propose that substrates principally modulate the kinetics, rather than the thermodynamics, of GadC’s energy landscape (Figure 8C).

Remarkably, the proposed conformational cycle deviates from those of structurally homologous sodium-coupled symporters, which have been shown to undergo substantial rearrangements in response to substrate binding^39–41,47^. However, our results may indicate that the contribution of substrates to the conformational equilibrium of GadC closely parallels the role played by the driving sodium ion in Mhp1 and vSGLT, as saturating concentrations had no effect on the thermodynamics of either symporter relative to ligand-free conditions^39,41,47^. Instead, sodium binding introduced an impassable kinetic barrier preventing uncoupled sodium flux down its concentration gradient^87,88^. We propose that glutamate and GABA play the inverse role in GadC: their binding reduces the kinetic barrier separating the inward- and outward-facing conformations while minimally affecting the underlying equilibrium. Similarly, their release into the cytoplasmic or periplasmic space introduces a large energy barrier to substrate free isomerization. We note that while the DEER data do not reflect changes in kinetic barrier, this proposal is consistent with previous results for the homologous serine-threonine antiporter SteT showing that ligand binding reduced a kinetic energy barrier during unfolding^89^.

A feature of our proposed conformational cycle is that the dynamics of TM10 distinguish between glutamate and GABA. Intriguingly, although AF2 predicted that this helix acts as an extracellular thin gate, our ensemble of models suggested that its motion was tightly coupled to that of TM9. In contrast, the DEER data reveal that TM10, but not TM9, distinguishes between glutamate and GABA binding. This was particularly striking given the observed correspondence between distance changes predicted by AF2 and distance changes observed by DEER (Figure S13). Further, the dynamics observed in TM10 closely mirrors the dynamics of its pseudosymmetric counterpart TM5 in poly-specific amino acid symporters, such as LeuT, the bending of which correlates with overall transport rate^90,91^. As GadC is highly specific for glutamate and GABA, it was not possible to establish an equivalent trend between transport rate and bending in TM10. We note that a similar mechanism of alternating access was experimentally observed in LAT1, a homologous broad-specificity amino acid exchanger^33,77^. Thus, whether this helix functions in a manner equivalent to TM5 in sodium-coupled amino acid symporters remains to be determined.

Our work presented here establishes a new experimental methodology for investigation of conformational changes underpinning transport. The ensembles of AF2 models, validated by DEER distance measurements, will allow further computational analysis, such as MD simulations, as well more detailed testing of transport mechanisms via site-directed mutants to trap specific conformations as has been shown for other transporters^92^.

## MATERIALS AND METHODS

### Site-directed mutagenesis

A codon-optimized version of the GadC gene from *Escherichia coli* str. O157:H7 (Genscript) was cloned into a pET19b vector encoding an N-terminal deca-histidine tag. A cysteine-less construct (C60V, C246A, C380V) was generated from this template using site-directed mutagenesis (QuikChange). All single- and double-cysteine mutants were similarly generated from this cysteine-free construct and verified by Sanger sequencing using both T7 forward and reverse primers.

### Expression, purification, and spin labeling of GadC

Plasmids encoding either wildtype or mutant GadC were transformed into competent *E. coli* str. C43 (DE3) cells and overexpressed in 1L minimal media A supplemented with ampicillin (Gold Biotechnology) as previously described. Upon reaching an absorbance (OD600) of 0.7-0.8, GadC expression was induced by adding 1 mM IPTG (Gold Biotechnology) and the temperature was dropped to 20°C. Cells were harvested after 16 hours by centrifugation at 5500 g for 15 minutes, resuspended in 22 mL lysis buffer (100 mM KPi, 10 mM DTT, pH 7.5), and lysed by sonication. After centrifugation at 9000 g for 15 minutes, the supernatant was collected and ultracentrifuged at 200,000 g for 90 minutes.

The pelleted membrane fractions were then solubilized in resuspension buffer (50 mM Tris/Mes, 200 mM NaCl, 20% glycerol, 1 mM DTT, pH 7.5) containing 1% β-DDM (Anatrace) and stirred on ice for 60minutes. Insoluble material was removed by ultracentrifugation at 200.000g for 30 minutes, and the supernatant was incubated with 1.0 mL Ni-NTA Superflow (Qiagen) resin at 4°C for two hours with 25 mM imidazole. After washing with ten column volumes of resuspension buffer containing 50 mM imidazole and 0.05% B-DDM, purified GadC was eluted from the resin using resuspension buffer with 250 mM imidazole and 0.05% β-DDM.

Following the addition of 60 mM Mes, single- and double-cysteine mutants were labeled with three rounds of 20-fold molar excess MTSSL (Enzo Life Sciences) per cysteine at room temperature and moved to ice overnight after four hours. Samples were then concentrated using Amicon Ultra 50.000 MWCO filter concentrators (Millipore) to a final concentration no greater than 3 mg/ml, as reported by absorbance at 280 nm (*ϵ*=67.840 M^−1^ cm^−1^), and purified into 200 mM Tris/Mes, pH 7.2, 20% glycerol, 0.05% β-DDM by size exclusion chromatography using a Shodex KW-803 column with guard column. Peak fractions eluted at 9.5-10.5 ml and were isolated for further studies.

### Reconstitution of GadC into proteoliposomes

A 3:1 ratio (weight/weight) of E. coli polar lipids and L-α-phosphocholine (Avanti Polar Lipids) were dissolved in chloroform and evaporated with a rotary evaporator. After overnight desiccation in a vacuum chamber, lipids were resuspended in the appropriate buffer, homogenized by ten cycles of freeze-thawing, and stored in small aliquots at -80°C.

Lipids prepared for liposomes were resuspended in 25 mM KPi, 150 mM KCl pH 5.5, and either 5 mM L-Glu or 5 mM GABA to a final concentration of 20 mg/ml (16.4 mM). Before reconstitution, lipids were diluted and destabilized with the addition of 1.25% octyl-β-D-glucopyranoside (β-OG) (Anatrace) and extruded through a 400 nm membrane filter (Whatman). Purified GadC was added to the sample at a 1:200 ratio (weight/weight), bringing the final lipid concentration to 5 mg/mL. Following a thirty-minute incubation at room temperature, detergent was removed from the sample by the gradual addition of 400 mg/mL SM-2 polystyrene Bio-Beads (Bio-Rad) over the course of four hours. After rocking overnight in the dark, the proteoliposome solution was cleared of biobeads and ultracentrifuged at 150,000 g for 60 minutes. Proteoliposomes were then resus-pended in external buffer (25 mM KPi, 150 mM KCl, pH 5.5) and ultracentrifuged to remove external substrates. After repeating this ultracentrifugation step a total of three times, proteoliposomes were suspended in external buffer at a final lipid concentration of 100 mg/ml. GadC concentration was then quantified using SDS/PAGE and densitometry (ImageJ v. 1.53g), with purified GadC in β-DDM serving as a standard curve.

### Transport assays

*In vitro* transport assays were carried out either in triplicate (concentration-dependent, Figures 1, S2, and S3) or in duplicate (time-dependent, Figure S1) as previously described^26^. An additional baseline measurement was performed on ice. Glutamic acid (between 25 μM and 1 mM) was added to external buffer and checked for pH immediately prior to all transport experiments. For the time-dependent transport Glu/GABA exchange assay shown in Figure S1, a fixed external Glu concentration of 50 μM at pH 5.5 was used. In both experiments, proteoliposomes (2 μL) were added to external buffer (98 μL) containing 1 μCi [^3^H]-L-glutamic acid (approximately 200 nM) and gently agitated. For titration experiments on wildtype GadC, proteoliposomes (1 μL) were added to external buffer (99 μL) containing 1 μCi [^3^H]-L-glutamic acid. Substrate uptake proceeded for two minutes at 25°C and was quenched by adding ice-cold stop buffer (25 mM glycine, 150 mM KCl, pH 9.5) and vacuum-filtering the solution through a 0.22 μm GSTF filter (Millipore) pre-soaked in stop buffer. The filter was then washed with an additional 6 mL stop buffer, removed, and added to 5 mL Ecoscint H scintillation solution (National Diagnostics). Following quantitation, data were analyzed using Michaelis-Menten kinetics using the *curve_fit* function implemented in SciPy^93^. Baseline measurements were subtracted from the 25°C measurements.

### Reconstitution of GadC into lipid nanodiscs

Lipids for nanodisc reconstitution were prepared as described above and resuspended in 50 mM Tris/Mes pH 7.5 to a final concentration of 20 mM. MSP1D1E3 was purified as previously described^94^. Nanodisc reconstitution proceeded using a molar ratio of 1:8 GadC:MSP1D1E3, 1:50 MSP1D1E3:lipid, and 1:5 lipid:cholate. Detergents were gradually removed from the solution using SM-2 Bio-Beads as previously described^94^. After overnight incubation, biobeads were removed from the solution using a 0.20 μm filter. Nanodisc-reconstituted GadC was then isolated from empty nanodiscs by size-exclusion chromatography using a Superdex 200 Increase 10/300 GL column into 50 mM Tris/Mes, pH 7.5, 10% glycerol and concentrated using an Amicon Ultra 100,000 MWCO filter concentrator (Millipore). The pH of all protein samples was carefully determined using a microelectrode and adjusted using 1 M citrate and 1 M Tris. Protein concentration was then evaluated using CW EPR spectroscopy as previously described ^95^. Glycerol was added to all DEER samples to a final concentration of 23% vol/vol, which were then flash-frozen in liquid nitrogen prior to DEER spectroscopy.

### CW-EPR and DEER spectroscopy and data analysis

Spin-labeled GadC was characterized using CW-EPR at 25°C using a Bruker EMX spectrometer operating at a frequency of 9.5 GHz, a 10 mW incident power, and a modulation amplitude of 1.6 G. DEER measurements were carried out using a dead-time free four-pulse protocol^96^ at either 50 K (for 143C/480C) or 83 K (all other double-cysteine mutants). Pulse lengths were as follows: 10 ns to 14 ns (first *π*/2 pulse), 20 ns (second and fourth *π* pulse), and 40 ns (third *π* pulse). The pump and observation frequencies were separated by 62.26 MHz. Echo decay data were analyzed into distance distributions using GLADDvu with the last 500 ns of the signal truncated. Fitting model parameters were chosen using the Bayesian Information Criterion^97^. To analyze the pH titration distance data collected using GadC 143C/480C, the long-distance component was isolated from the two short-distance components, and was fitted with a sigmoid function using the *curve_fit* function as implemented in SciPy^93^. For all DEER pairs, the distance distributions were compared to predictions generated by MDDS, which was accessed using the CHARMM-GUI web server^69^.

### Generation of structural models in multiple conformations using AlphaFold2 and Rosetta

The structure of GadC was modeled using AlphaFold v.2.0.1 using a modified version of ColabFold^64,98^. Multiple sequence alignments were generated using MMSeqs2^99^. Several modifications were introduced to the default pipeline to obtain alternative conformations of GadC^65^. First, all inward-open structures were removed from the list of templates fetched by MMSeqs2 prior to modeling, leaving inward-facing occluded structures of ApcT^78^ (PDB 3GIAa and 3GI9c) and GkPApcT^100^ (PDB 5OQTa and 6F34a), as well as outward-facing structures of AdiC^71,72^ (PDB 3OB6b and 5J4Ib) and LAT1^33,77^ (PDB 7DSQb). To ensure that all templates were used, the sequence identity cut-off for inclusion was lowered from 10% to 1%. Second, the configuration *subsample_templates*, which is set to *False* by default, was set to *True*. Third, the depth of the MSA (set by *max_msa_clusters*) was reduced to between eight and 20 sequences, and the total number of sequences provided to the protocol (*max_extra_msa*) was set to double that number. Finally, refinement by OpenMM was replaced by Cartesian minimization using Rosetta FastRelax^75^. For this step, the *membrane_highres_Menv_smooth* scoring function was used, with the score terms *coordinate_constraint* set to 1.0, *cart_bonded* set to 0.5, and *pro_close* set to 0.0^74^. The membrane bilayer’s position was calculated using OCTOPUS as previously described^101^. The Cartesian coordinates of all backbone heavy atoms were constrained to their initial positions by weights inversely proportional to their predicted local distance difference test (pLDDT^102^) values:

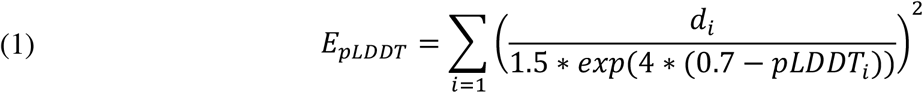

Here, *d*_*i*_ is the *i*th atom’s distance from its initially predicted position, and *E* refers to the coordinate constraint energy term^103^ (we note that the pLDDT values provided by AF2 range from 0 to 100 and were therefore divided by 100 prior to calculating these values). The side chain heavy atoms of residues with pLDDT values exceeding 0.9 were similarly constrained in Cartesian space. This pipeline was used to generate 650 structural models of GadC without its C-terminal domain (residues 471-511), with twenty-five models generated for each MSA depth by each of the two AF2 neural networks capable of using templates. These models were then aligned to the crystal structure using TM-Align^104,105^ and projected onto a lower-dimensional space using principal component analysis (PCA) as implemented by SciKit-Learn^106^. Only the positions of alpha carbons belonging to helical residues were considered, and loop residues were omitted from this analysis.

## ACKNOWLEDGEMENTS

This study was funded by the National Institutes of Health (GM 128087). We would like to thank Dr. Derek P. Claxton for critical review of this manuscript and Drs. Richard Stein and Eric J. Hustedt for fruitful discussions regarding the interpretation of DEER confidence bands.

## SUPPLEMENTAL FIGURES

**Supplemental Figure S1.**
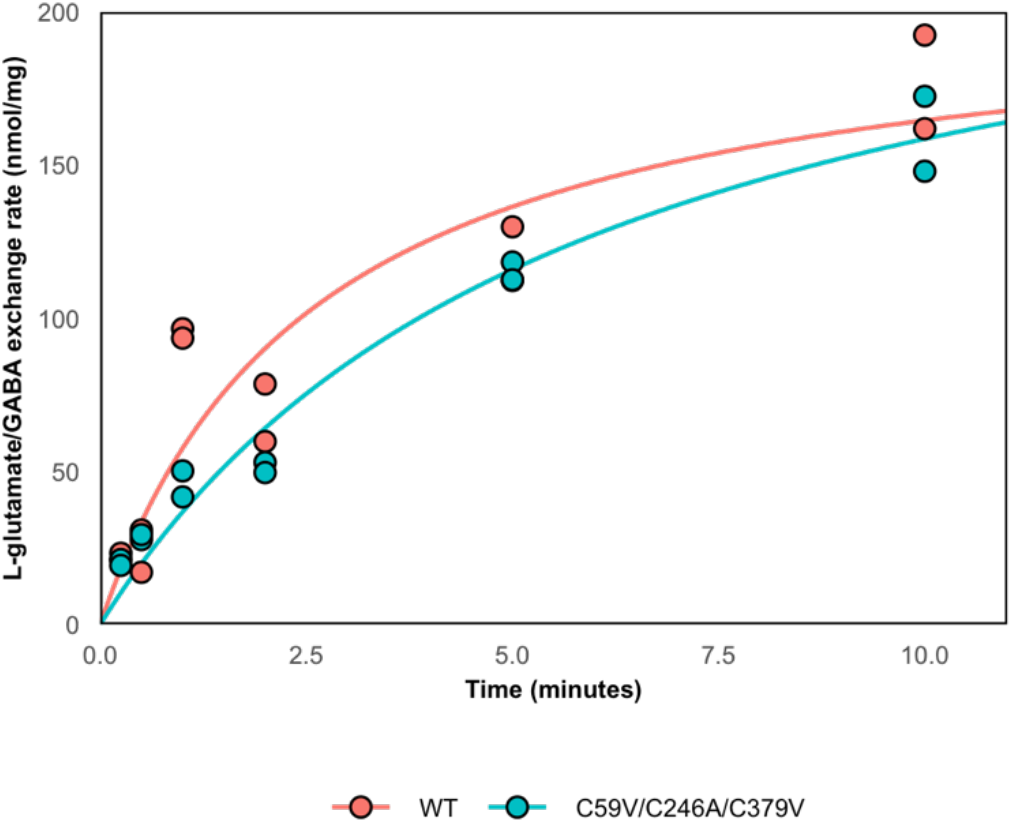
Time-dependent glutamate/GABA antiport in either wildtype or cysteine-less construct of GadC (C59V/C246A/C379V) reconstituted into proteoliposomes.

**Supplemental Figure S2.**
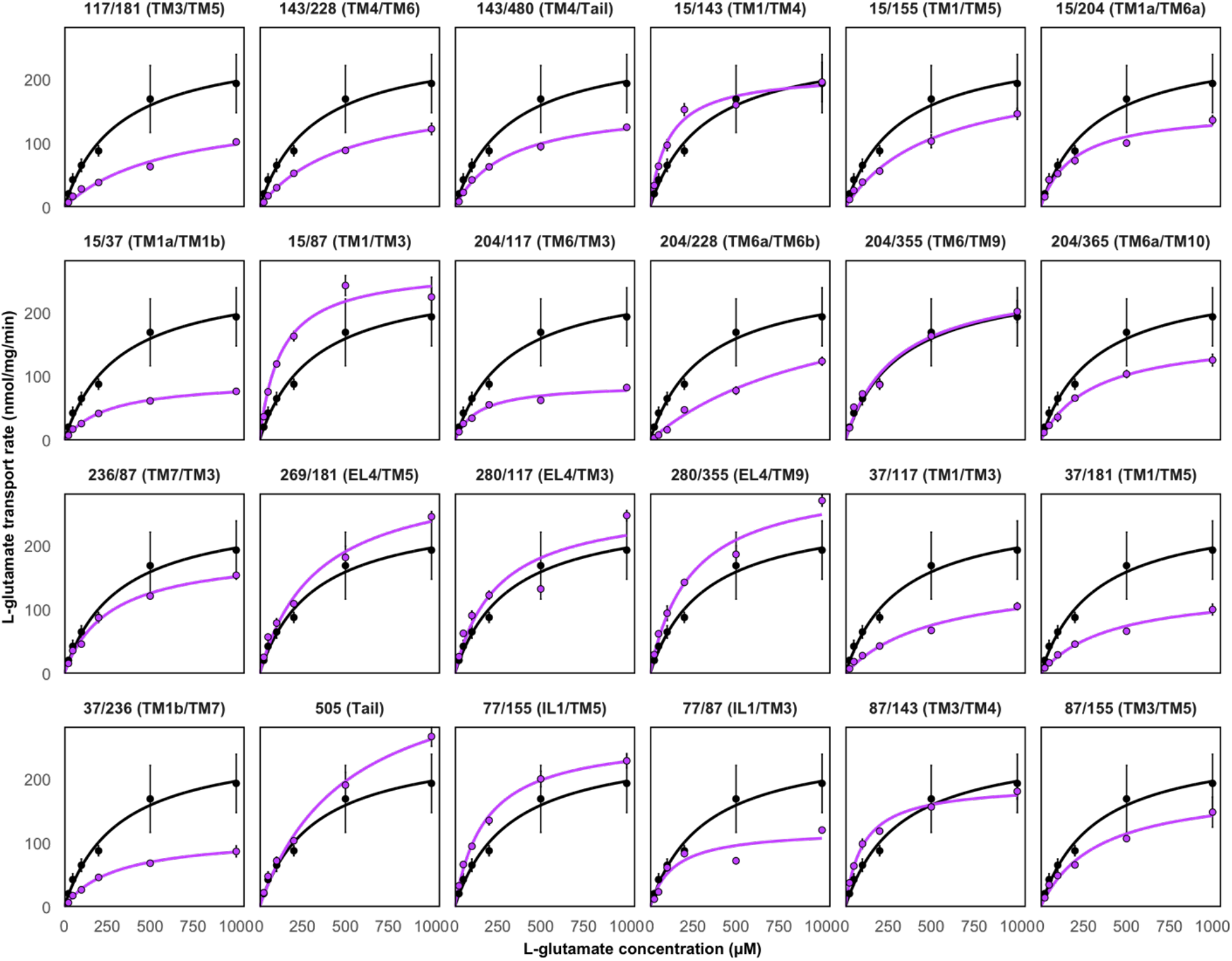
Glutamate transport at pH 5.5 of double-cysteine GadC mutants in proteoliposomes following spin labelling. WT activity shown in black for comparison, while mutant curves are shown in purple. Error bars correspond to the standard error of the mean.

**Supplemental Figure S3.**
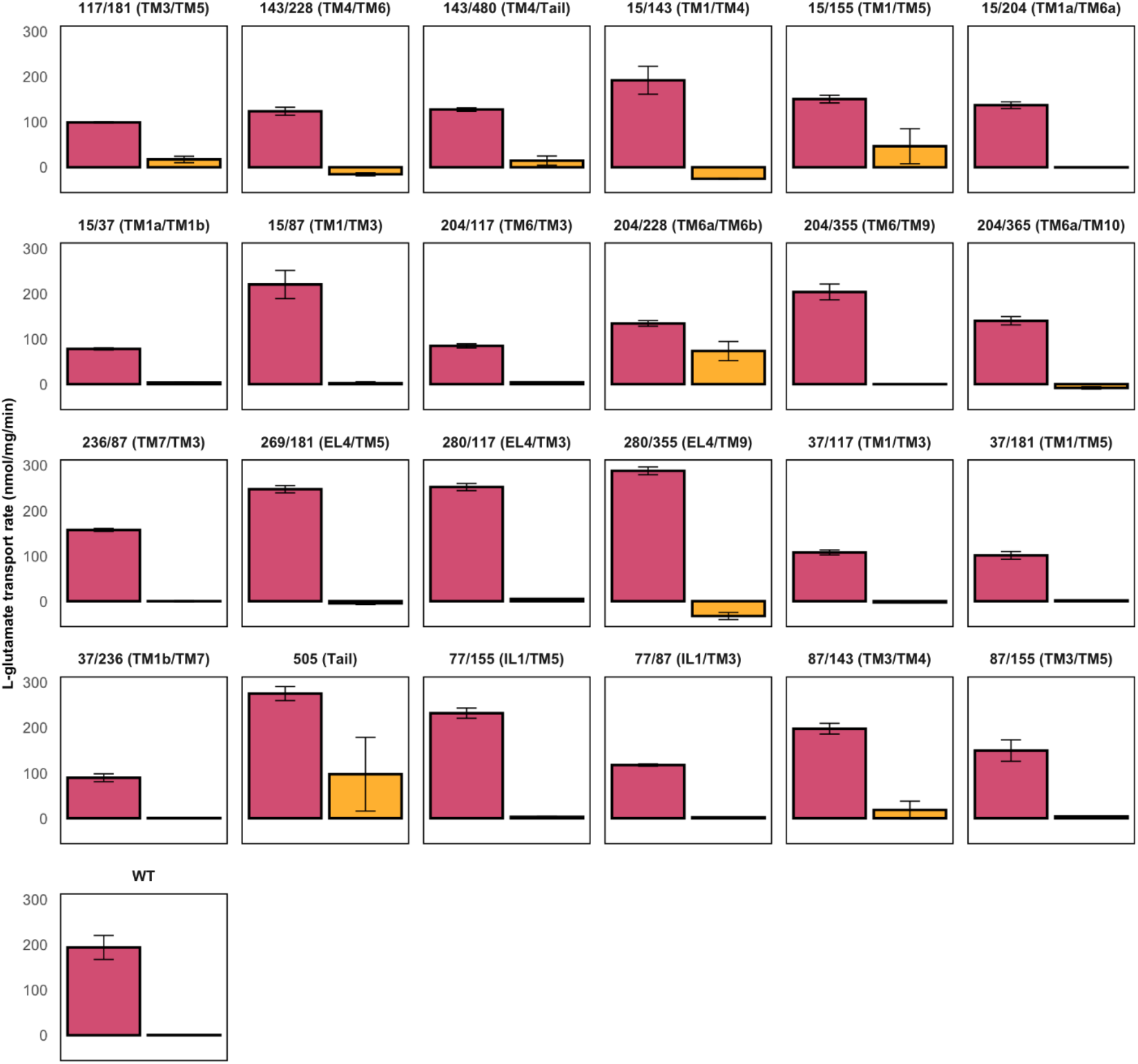
Glutamate transport of double-cysteine GadC mutants in proteoliposomes following spin labelling at pH 5.5 (red) and 7.5 (yellow). Error bars correspond to the standard error of the mean.

**Supplemental Figure S4.**
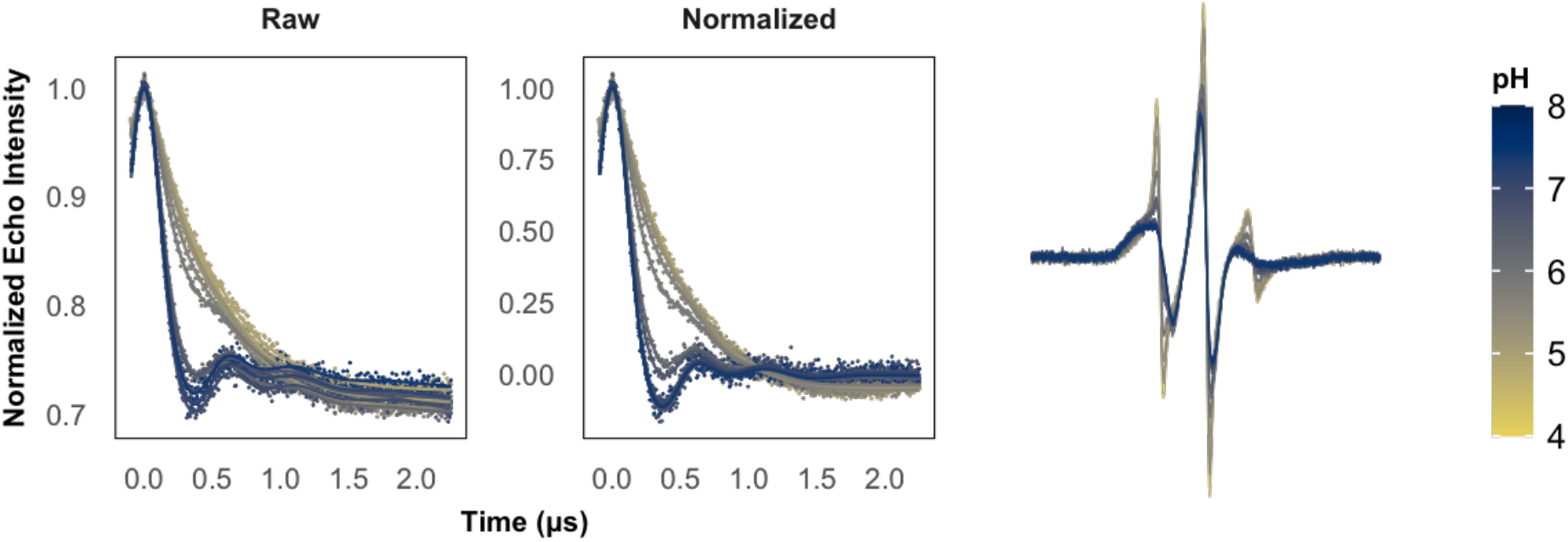
pH-dependent DEER data and continuous-wave EPR spectra of GadC 143/480.

**Supplemental Figure S5.**
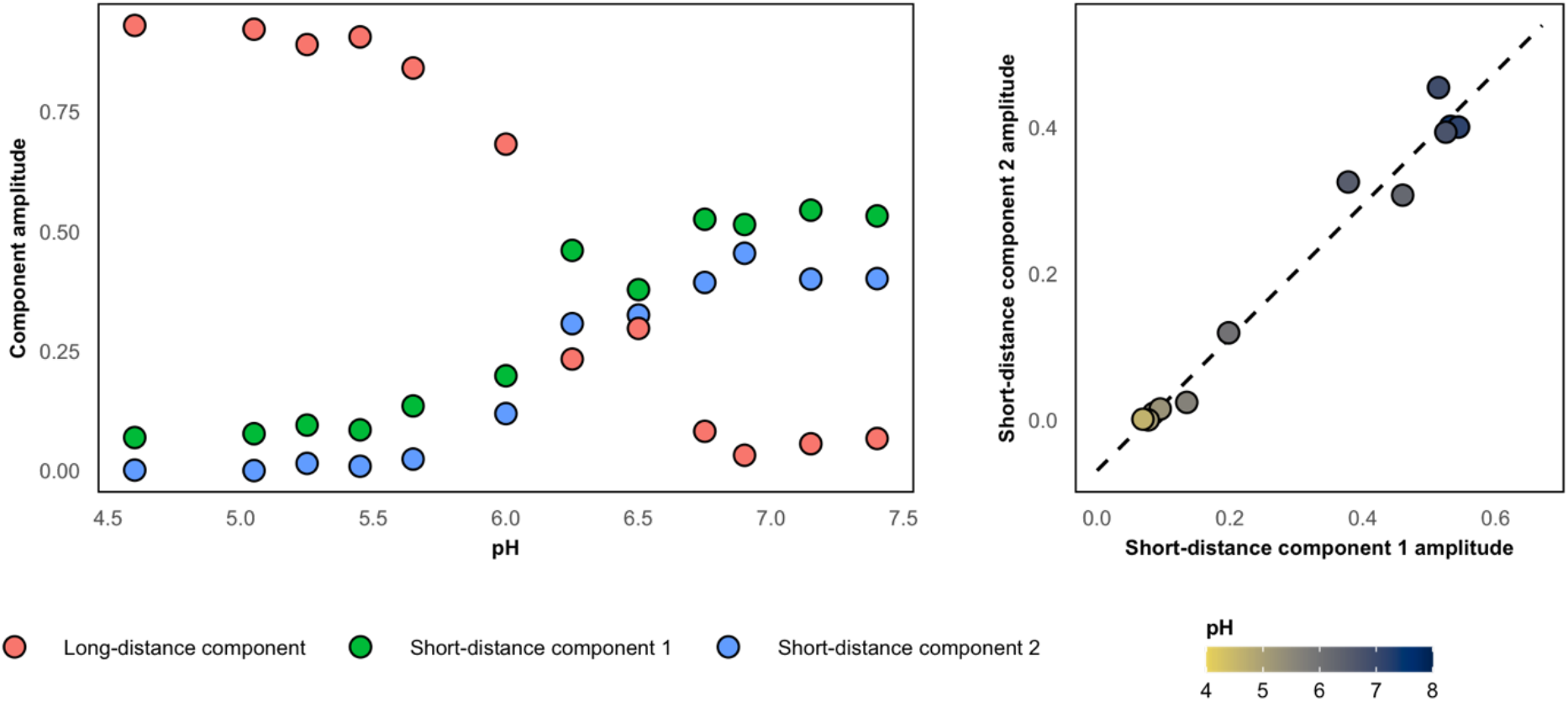
Correlation between short-distance DEER components in spin pair 143/480 during detachment of the C-terminal tail.

**Supplemental Figure S6.**
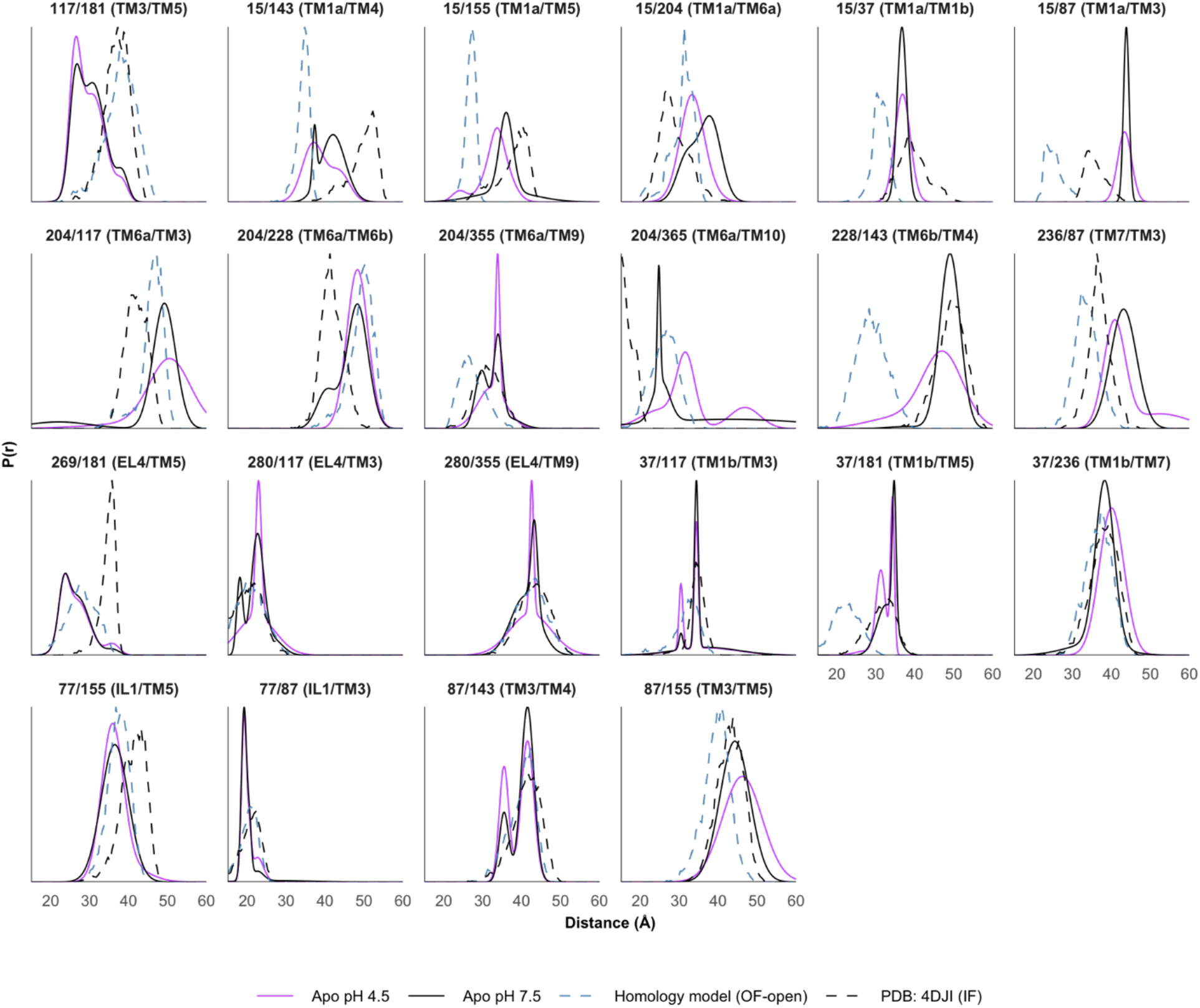
Comparison of experimental DEER data to predictions made from an outward-facing homology model generated using Rosetta. Deviations were primarily observed on the extracellular pairs and on TM1 on the intracellular side.

**Supplemental Figure S7.**
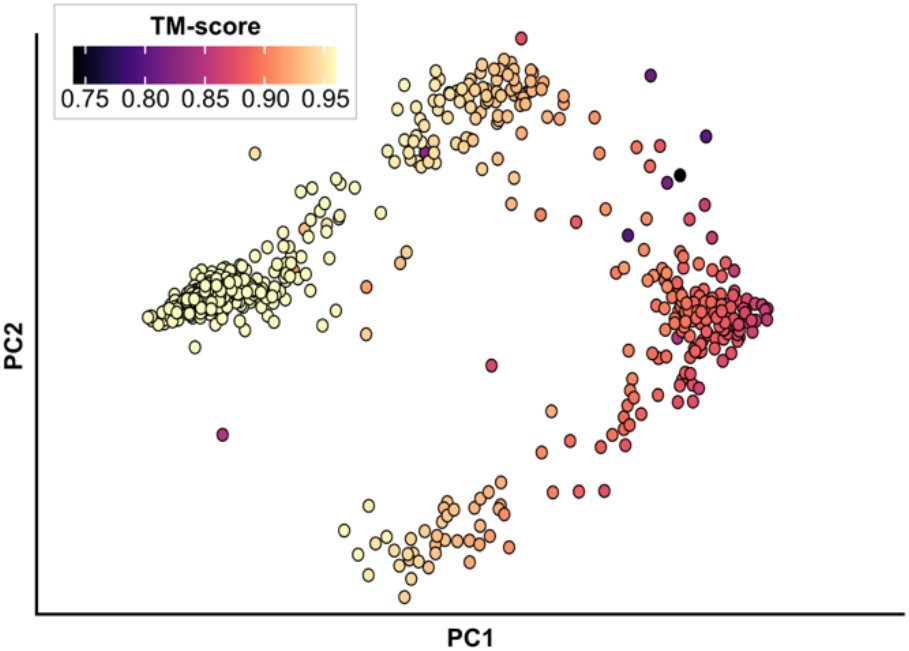
Similarity of AlphaFold2 models to inward-facing crystal structure. Similarity was evaluated using TM-score, which ranges from 0 to 1 with greater values indicating greater similarity.

**Supplemental Figure S8.**
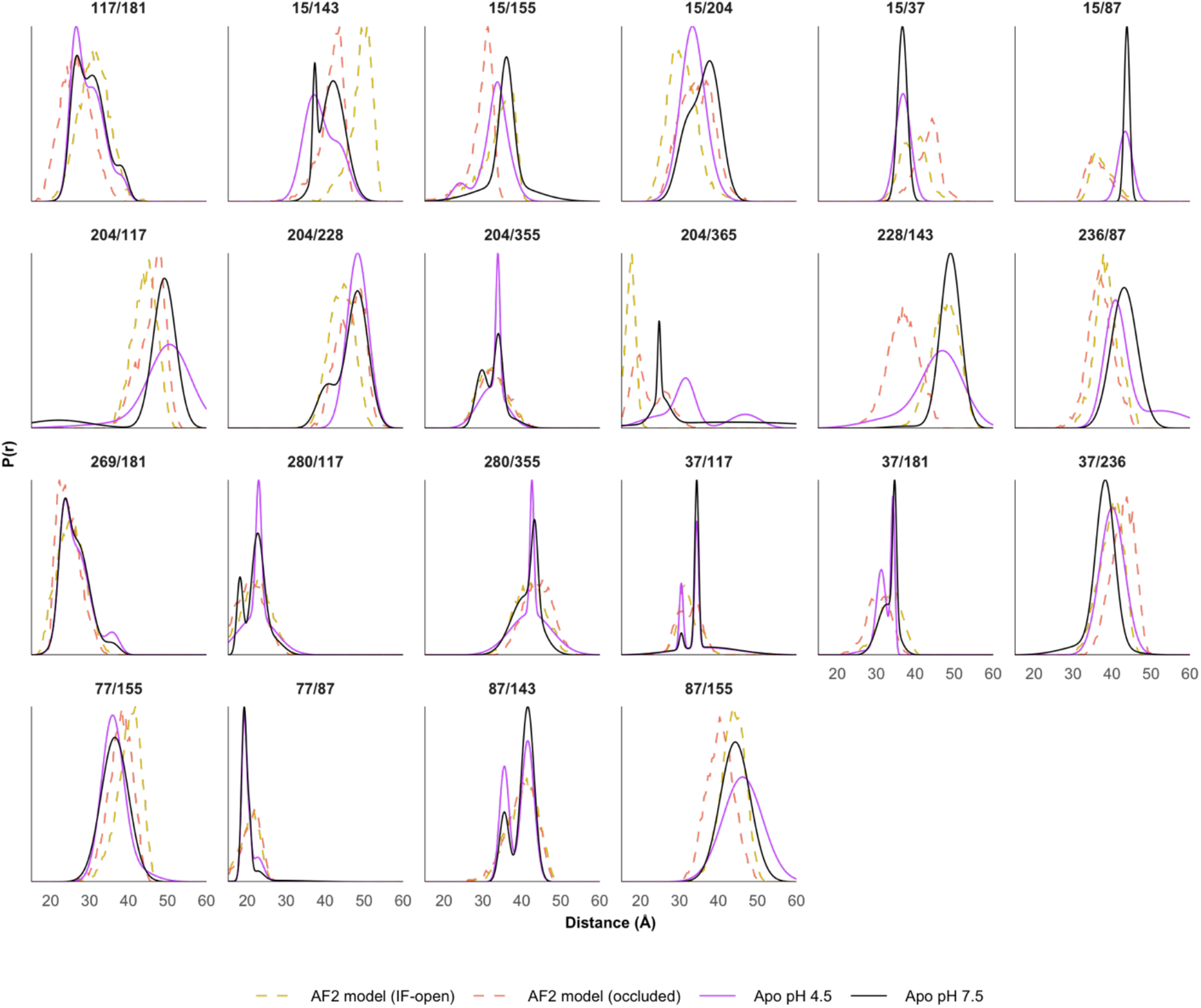
Comparison of experimental DEER data to predictions made from inward-open (gold) and fully occluded (orange) models generated using AlphaFold2.

**Supplemental Figure 9.**
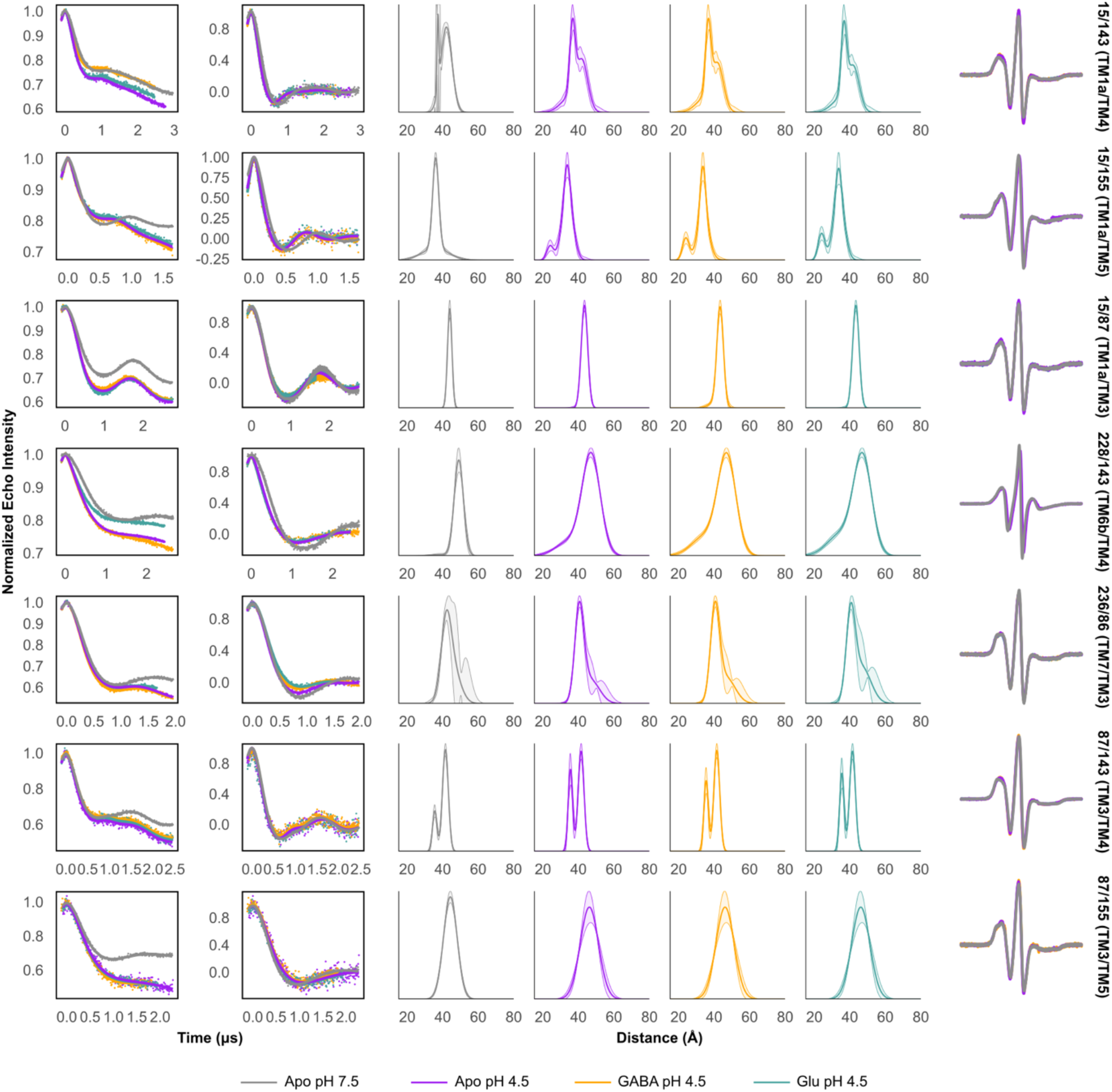
Experimental data for spin pairs involved with evaluating interdomain movements on the intracellular side. From left: Raw DEER data, background-corrected DEER data, distance distributions with 95% confidence intervals calculated using GLADDvu, and continuous wave EPR spectra.

**Supplemental Figure 10.**
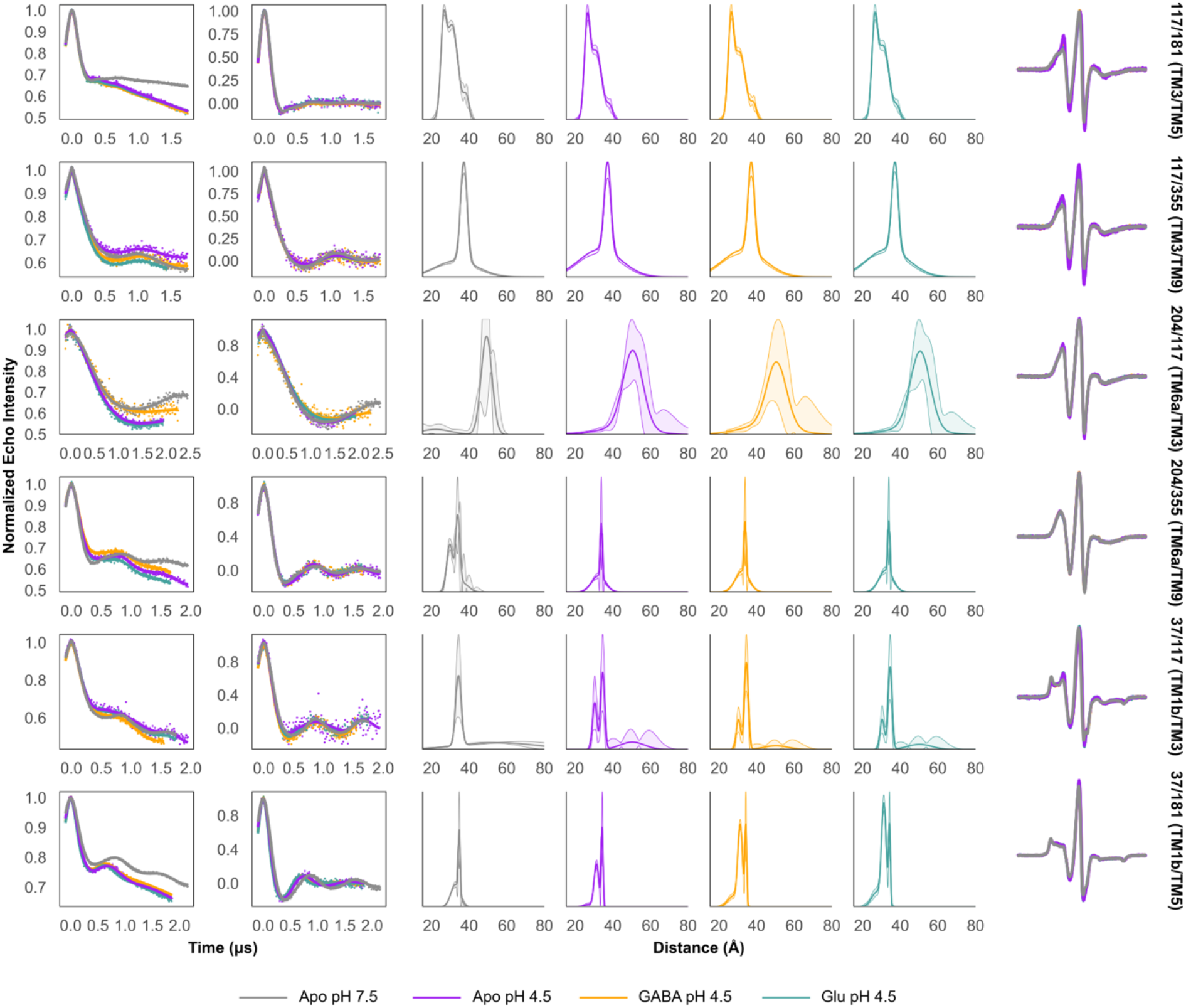
Experimental data for spin pairs involved with evaluating interdomain movements on the extracellular side. From left: Raw DEER data, background-corrected DEER data, distance distributions with 95% confidence intervals calculated using GLADDvu, and continuous wave EPR spectra.

**Supplemental Figure 11.**
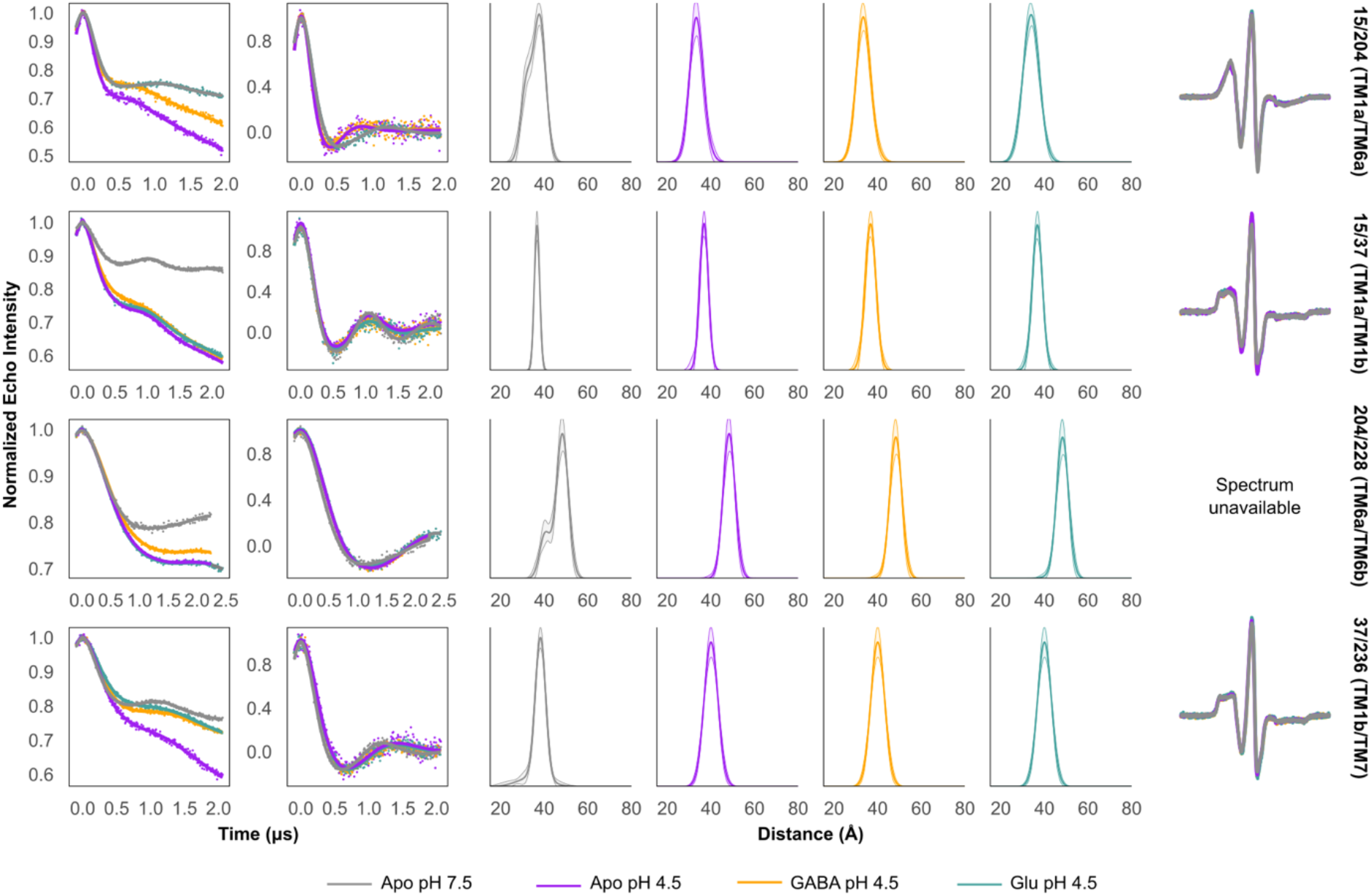
Experimental data for spin pairs involved with evaluating intradomain movements in the bundle domain. From left: Raw DEER data, background-corrected DEER data, distance distributions with 95% confidence intervals calculated using GLADDvu, and continuous wave EPR spectra.

**Supplemental Figure S12.**
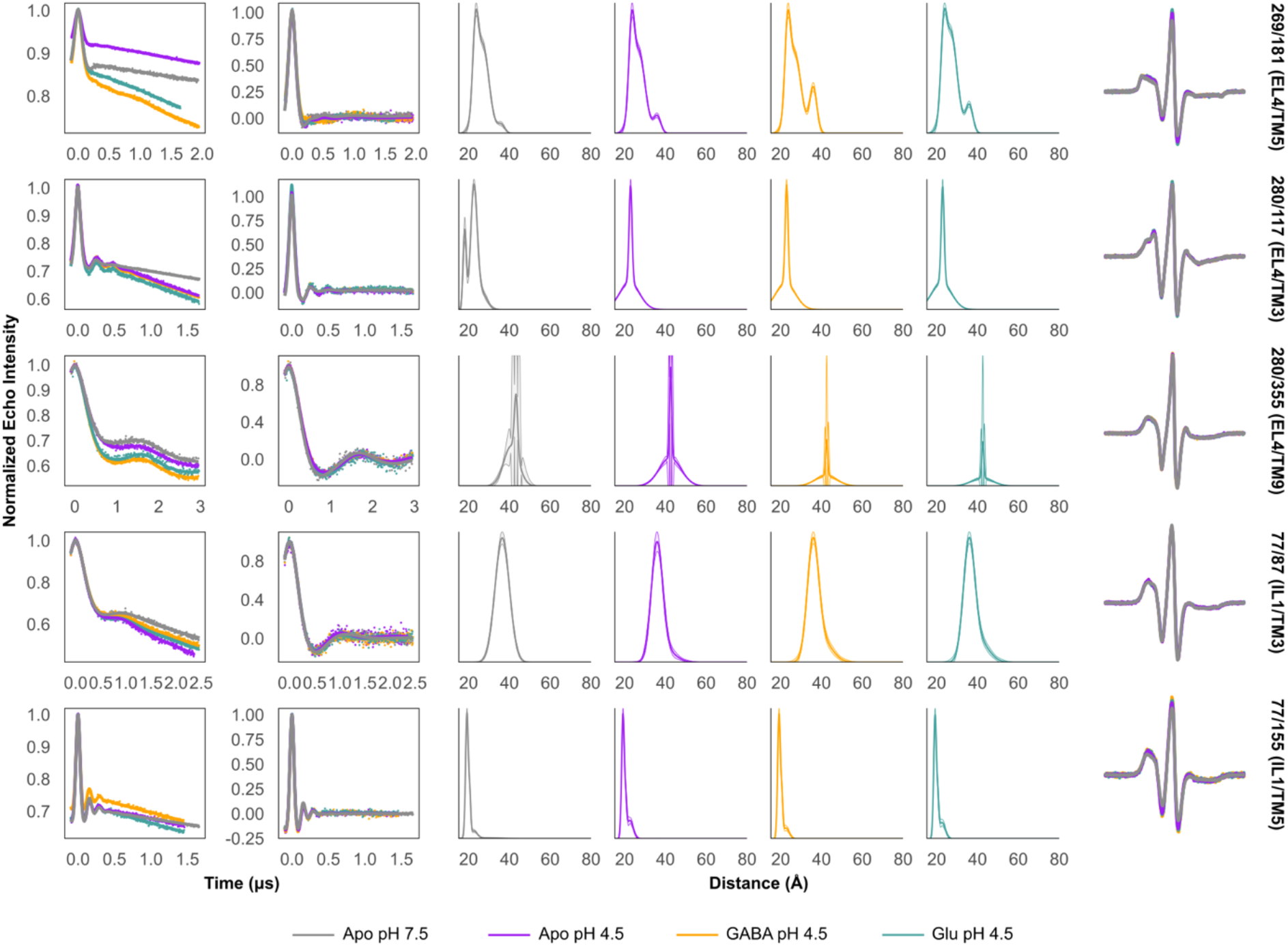
Experimental data for spin pairs involved with evaluating movements of EL4 and IL1. From left: Raw DEER data, background-corrected DEER data, distance distributions with 95% confidence intervals calculated using GLADDvu, and continuous wave EPR spectra.

**Supplemental Figure S13.**
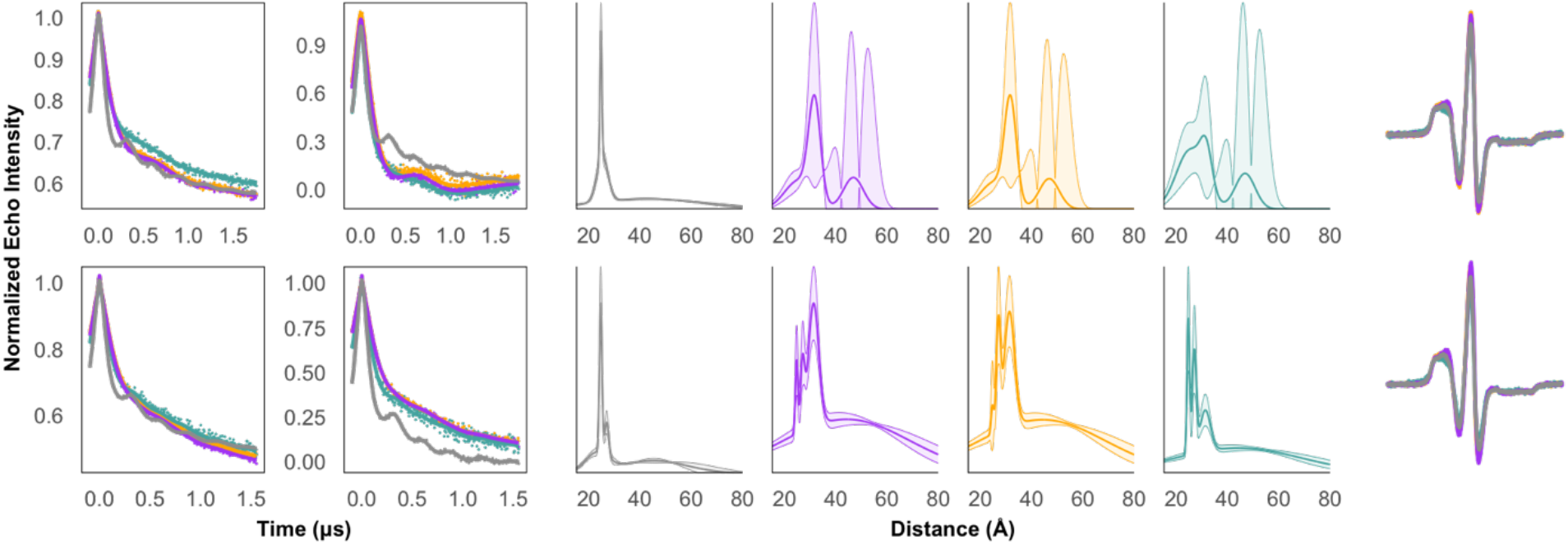
Experimental data and biological repeats of 204/365. From left: Raw DEER data, background-corrected DEER data, distance distributions with 95% confidence intervals calculated using GLADDvu, and continuous wave EPR spectra.

**Supplemental Figure S14.**
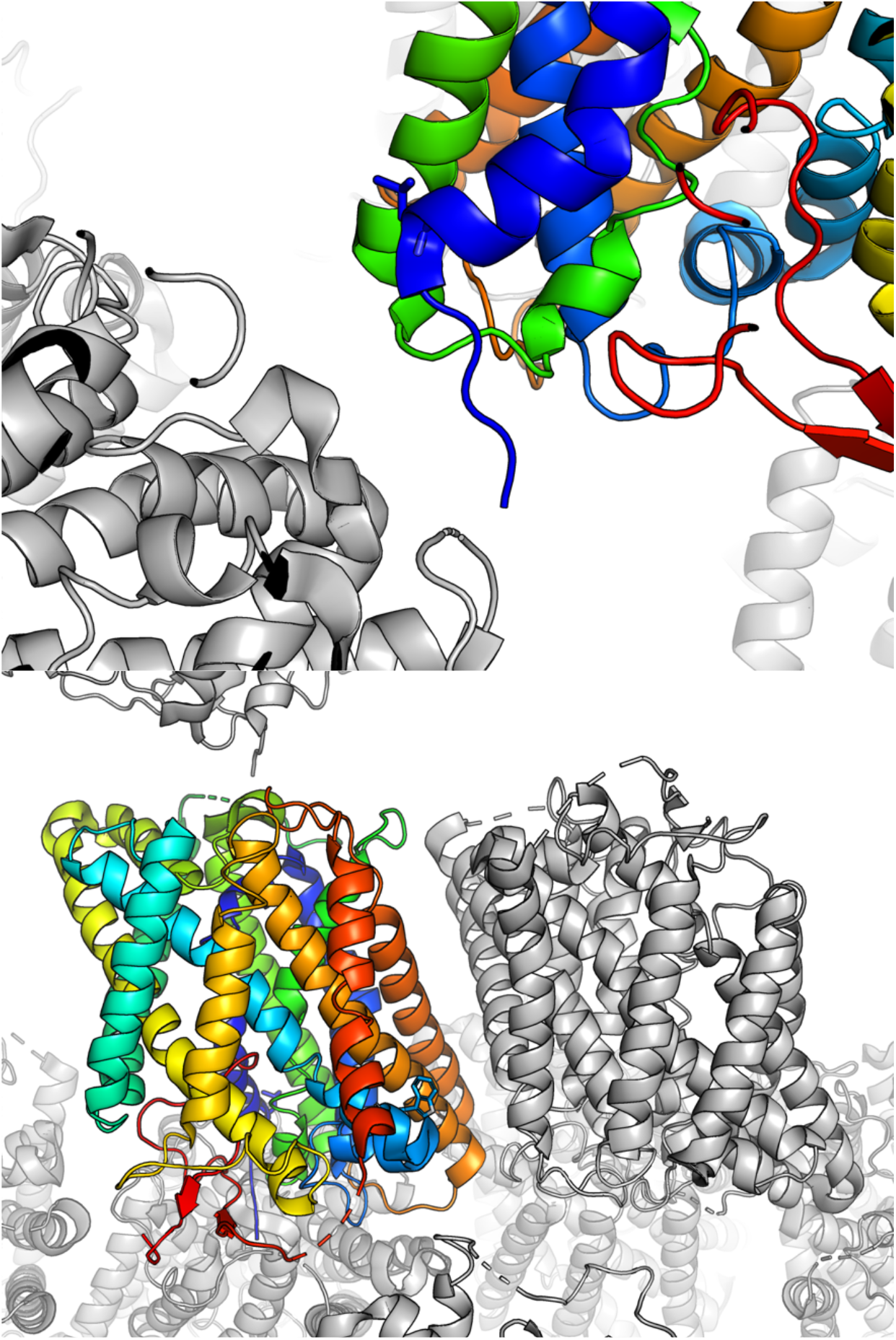
Crystal contacts near residues 15 and 87 may contribute to deviations between predicted and experimental DEER distance distributions. **Top:** residue L15 shown in dark blue on TM1a. The nine N-terminal residues of GadC were not resolved. **Bottom:** residue W87 shown in light blue. Crystallographic symmetry was calculated using PyMol.

**Supplemental Figure S15.**
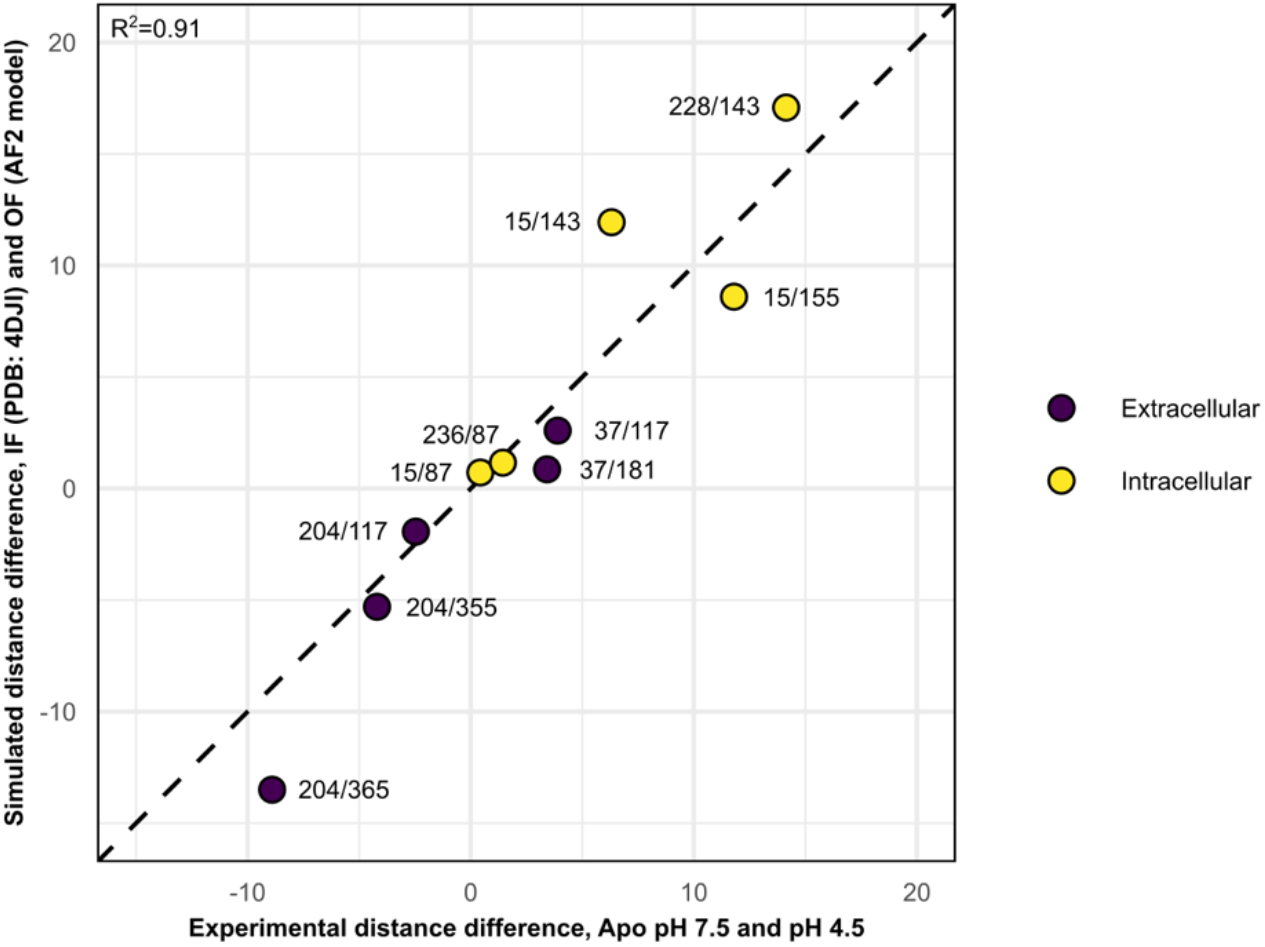
Correlation between experimental distance changes observed using DEER and predicted distance changes between the inward-facing experimental structure and the outward-facing AF2 (AF2) model. This plot is limited to pairs with spin-labeled residues positioned on helices predicted to undergo large-amplitude motions.

